# GWAS from Spoken Phenotypic Descriptions: A Proof of Concept from Maize Field Studies

**DOI:** 10.1101/2023.12.11.570820

**Authors:** Colleen F. Yanarella, Leila Fattel, Carolyn J. Lawrence-Dill

## Abstract

We present a novel approach to Genome-Wide Association Studies (GWAS) by leveraging unstructured, spoken phenotypic descriptions to identify genomic regions associated with maize traits. Utilizing the Wisconsin Diversity panel, we collected spoken descriptions of *Zea mays* ssp. *mays* traits, converting these qualitative observations into quantitative data amenable to GWAS analysis. First, we determined that visually striking phenotypes could be detected from unstructrured spoken phenotypic descriptions. Next, we developed two methods to process the same descriptions to derive the trait plant height, a well-characterized phenotypic feature in maize: (1) a semantic similarity metric that assigns a score based on the resemblance of each observation to the concept of ‘tallness,’ and (2) a manual scoring system that categorizes and assigns values to phrases related to plant height. Our analysis successfully corroborated known genomic associations and uncovered novel candidate genes potentially linked to plant height. Some of these genes are associated with gene ontology terms that suggest a plausible involvement in determining plant stature. This proof-of-concept demonstrates the viability of spoken phenotypic descriptions in GWAS and introduces a scalable framework for incorporating unstructured language data into genetic association studies. This methodology has the potential not only to enrich the phenotypic data used in GWAS and to enhance the discovery of genetic elements linked to complex traits, but also to expand the repertoire of phenotype data collection methods available for use in the field environment.

## 1 INTRODUCTION

Collecting phenotype data can be slow, which limits the speed of association genetics and genomic studies for trait improvement. High-throughput phenotyping methods are an area of development that concentrates on engineering sensors and unmanned vehicles to collect (mainly visual) data about traits of various crop species (reviewed in Yang et al., 2020). These methods are beneficial for collecting large amounts of data in an automated fashion, but there are difficulties in deploying these tools in a field environment, and some traits are not detectable by images alone. Additionally, manually collecting phenotypes with pen-and-paper or tablet- and-stylus is time-consuming and generally requires predefined traits of interest. Sensors, imaging, and barcodes make data organization easier for large quantities of data (Yao et al., 2021; Sarić et al., 2022; Kazic, 2020). An underdeveloped area of in-field phenotyping ripe for exploration is using natural language descriptions of plants. Platforms exist where audio descriptions are recorded (Kazic, 2020). However, the biologically relevant data in spoken phenotypes thus far has remained inaccessible for association studies and other applications.

As demonstrated by Oellrich, Walls, and colleagues, language-based descriptions of phenotypes can be analyzed computationally to recover known biological associations in plants (2015). Their work used phenotype descriptions derived from various ontologies, then structured the data as entity quality (EQ) statements (where an entity is a feature, e.g., whole plant, and quality is a describer, e.g., dwarf-like), which results in less intensive computations (Mungall et al., 2010). Semantic or word-meaning similarity methods have also shown promise in ascertaining biologically meaningful genetic associations without the burden of manual curation (Braun et al., 2020; Braun and Lawrence-Dill, 2020). Additionally, pretrained models have enabled free-text descriptions of plant phenotypes for association studies based on semantics (Braun et al., 2021). The developments in the computational processing of natural language plant phenotypes through unstructured text formats contribute to conceptualizing methods for recording spoken descriptions of phenotypes.

Given the various difficulties associated with collecting structured phenotype data for trait analyses alongside recent advances in semantic reasoning by computer systems, we were curious to find out whether computing on unstructured descriptions of phenotype might now be tractable for biological inferences in a field setting. Imagine collecting phenotype data simply by walking through a field and using spoken words and vocabularies to describe phenotypes, then taking those audio recordings back to the lab as input along with genomics data to run genome-wide association studies. Are the analytical tools advanced enough for this kind of approach to phenotypic data collection and analysis? This question is the basis for experiments and results presented here.

To test whether associations between genomic variants and phenotypes could be derived from unstructured spoken language, we reasoned that a well-characterized diversity panel that had many associations already derived would be required as ground truth, which led us to select the Wisconsin Diversity (WiDiv) panel for this work. WiDiv (1) was developed to grow in the upper midwestern region of the United States; (2) includes only lines with similar flowering time and phenology, and (3) is made up of lines that have both genetic and phenotypic diversity (Hansey et al., 2011). Researchers using the WiDiv panel have increased the included accessions, investigated flowering time and biomass yield traits, and generated genetic marker data for the panel (Hansey et al., 2011; Hirsch et al., 2014; Mazaheri et al., 2019; Mural et al., 2022b).

Because the WiDiv panel data has expansive genotypic and phenotypic trait data available, we collected spoken descriptions of phenotypes for numerous traits (height, leaf width, stalk, tassel, leaf, braceroot, and overall color, etc.) during the summer of 2021 (Yanarella et al., 2023a). The objectives of this research were to (1) detect phenotype descriptions from spoken language, (2) demonstrate techniques for extracting phenotype data for a proof of concept analysis involving the plant height trait for GWAS, (3) perform GWAS with phenotypes derived from speech, and (4) use available gene function data to review and assess known and novel gene trait associations.

The methodologies outlined in this paper are versatile enough to analyze a range of traits present in the datasets; however, our demonstration centered on plant height due to its well-established genetic associations. It is important to acknowledge the subjective element introduced by using spoken, natural language descriptions, which diverges from the strict reproducibility and comparability offered by measured phenotypes and controlled vocabularies. Despite this, the ease of data gathering could prove beneficial, especially when it leads to the discovery of previously unknown gene-trait associations. Our successful identification of known genetic associations to the plant height trait validates this novel approach and sets a precedent for the future of phenotypic data acquisition and analysis. This study envisions a future where one can simply walk into a field, articulate observations, and receive trait-gene associations in return, leveraging the natural language as a powerful analytical tool.

## 2 MATERIALS AND METHODS

We used a genotypic dataset that includes WiDiv panel taxa (lines). A dataset of 18 million SNP markers (Mural et al., 2022a) obtained from aggregating RNA-Seq and resequencing techniques for 1,051 taxa (described in Mural et al. 2022b). Phenotypic datasets (described in Yanarella et al. 2024), include audio recordings as well as manual measurements of plant height for 686 unique WiDiv panel taxa (Yanarella et al., 2023a), hereafter referred to as the Yanarella *et al*. dataset. This dataset contains an additional 25 taxa (Supplementary Table 1) that were positive controls for the analysis of spoken descriptions of phenotypes, as these genetics lines were expected to have noticeable and readily describable phenotypes. These positive control taxa are not members of the WiDiv panel. Informed consent for the spoken data from participants was collected per Iowa State University’s Institutional Review Board’s (IRB) Exempt Project status, and volunteers provided informed consent for using their spoken observations. Phenotypic data include measurements for plant height and spoken descriptions of plants grown in a field environment. The Mural *et al*. and Yanarella *et al*. datasets share 653 taxa (Figure 1), which were used for further analyses described here.

**Figure 1.**
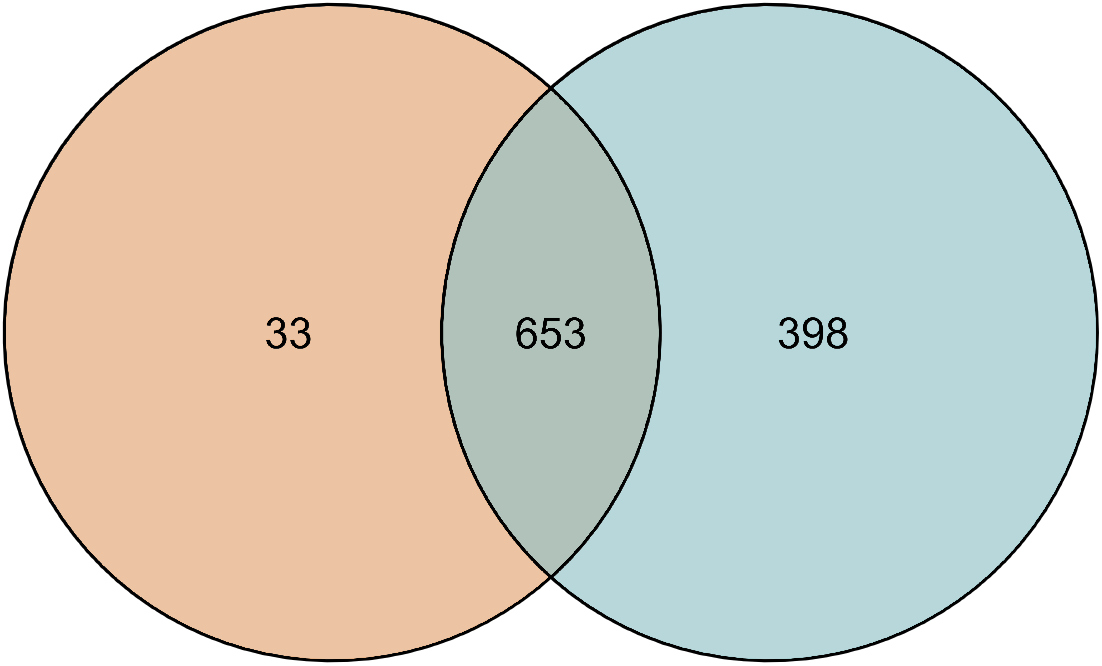
Comparison of intersections of Mural *et al*. and Yanarella *et al*. WiDiv dataset taxa (positive controls not included), where n is the number of unique taxa in each dataset. Mural *et al*. dataset (in blue, n = 1,051), and Yanarella *et al*. dataset (in orange, n = 686).

### 2.1 Spoken Phenotype Collection Summary

We reported on phenotype descriptions recorded by de-identified student workers as a peer-reviewed dataset elsewhere (Yanarella et al., 2024). We refer to the Yanarella *et al*. dataset throughout, and detailed information on data collection and dataset formats can be accessed via Yanarella *et al*. (2024). These data and the methods used for their collection are reviewed in brief here for context.

Unstructured spoken descriptions were exclusively captured in the field environment. The field in which the recordings were taken included two replicates planted in a randomized incomplete block design. The first block consisted of 31 WiDiv panel taxa for a seed increase, and the second block consisted of 8 B73 experimental control rows, 25 positive control taxa, and 655 unique WiDiv panel taxa. The second block contained two rows of the WiDiv panel line MEF156-55-2. Therefore, the recordings were taken over 720 rows in each replicate. Manually collected phenotypes included plant height as well as ear and tassel features. Phenotype data were collected during July and August of 2021, a time when plants had reached maturity.

Each of the de-identified student workers selected NATO code names. The students who were undergraduate Agronomy, Biology, and Genetics Majors at Iowa State University are known as “Delta”, “Golf”, “Kilo”, “Lima”, “Mike”, “Quebec”, “Victor”, “Yankee”, and “Zulu”. Each participant was instructed to state their NATO code name and the row tag number before observing the plants in each row (Figure 2a). This procedure ensured the participant’s de-identified connection to the row number and spoken observation while enabling the parsing of each observation so that multiple row observations could be recorded in the same file (data available from Yanarella et al., 2023a, and described in Yanarella et al., 2024). Participants were asked to comment on overall appearance, including color and height, tassel, ear, leaves, braceroots, and any disease. Two examples of how a given row might be described are as follows. “Alpha, 1641. Plants are flowering, there’s a lot of variability in height, tassels have emerged, the anthers have everted, brace roots at two nodes, some braceroots point upward.” Another: “Alpha, 2287. Plants are tall. They have brace roots at up to three nodes. Those brace roots are chunky. They have red to purple anthocyanins in rings. Leaves are very much upright. Silks are yellow, tassels are yellow.” Because many individuals passed each row many times, each row has a diversity of descriptions in the full dataset.

**Figure 2.**
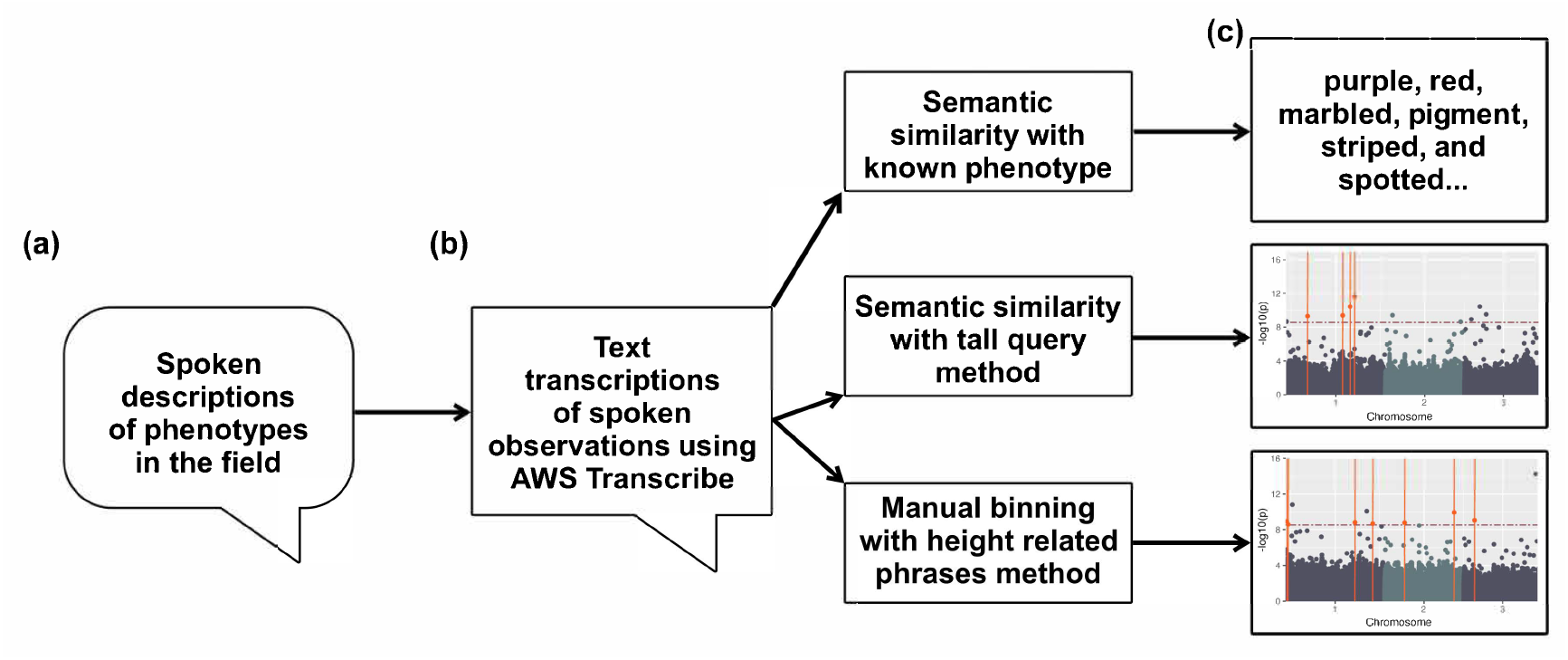
Spoken phenotype process overview. (a) In field spoken phenotype descriptions collection, (b) Spoken phenotype data processing, including transcript production and methods for generating numeric representations of phenotypes for traits, and (c) Top box: detecting phenotypes from positive control accession descriptions. Lower two boxes: GWAS using data derived from spoken observations.

### 2.2 Phenotype Detection from Spoken Descriptions of Positive Control Lines

A subset of spoken observation transcripts collected in the field containing the positive control accessions were parsed. Four to six terms from the description and phenotype records for each accession were drawn from MaizeGDB (Woodhouse et al., 2021). One term from these lists was used to collect synonyms from Merriam-Webster (Merriam-Webster, 2023) and WordHippo (Kat IP Pty Ltd, 2008) thesaurus services (Table 1). The number of rows containing at least one synonym related to each accession’s descriptions and phenotype records was calculated as a proportion to the number of observations for that accession.

**Table 1.**
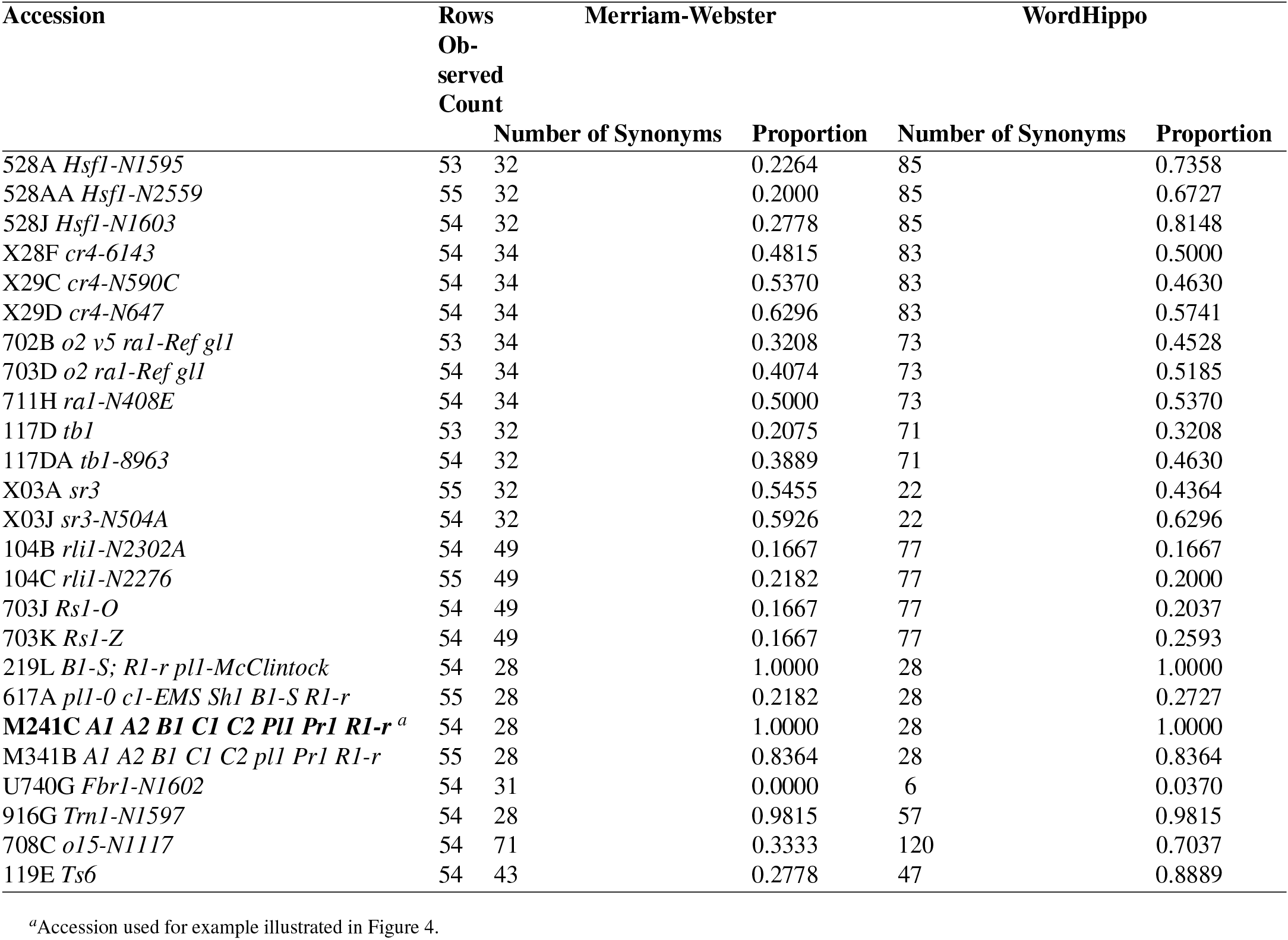
Proportion of word usage for describing positive control accessions.

### 2.3 Preprocessing Phenotypic Datasets

#### 2.3.1 Plant Height Measurement Data

R Scripts (v.4.2.2 and v.4.3.1) (R Core Team, 2023) were developed to process the measuring and scoring data from Yanarella *et al*. dataset such that only plant height observations were retained for each of the three observation sets (each row in the field was measured and scored by two teams made up of participants and one observation set was collected by a volunteer). Replicate number was programmatically added to these data, and positive controls were removed. The 653 taxa shared between the datasets were retained.

Best Linear Unbiased Estimators (BLUE) values were calculated for each taxon using R’s built-in lm function to perform linear regressions and the emmeans v.1.8.7 package (Lenth, 2023), where taxa and replicate were fit as fixed effects. Best Linear Unbiased Prediction (BLUP) corrected mean values were calculated for each taxon with the lmer function of the lme4 v.1.1-34 package (Bates et al., 2015) where taxa, replicate, and row number were fit as random effects. Visualization of diagnostics plots of the models (Supplementary Figure 1, Supplementary Figure 2) were generated using the ggResidpanel v.0.3.0 package (Goode and Rey, 2022).

#### 2.3.2 Semantic Similarity for Plant Height Spoken Data

Text transcripts of spoken data from the Yanarella *et al*. dataset were processed using Python v.3.8.2 (Van Rossum and Drake, 2009). The spaCy v.3.5.1 package (Honnibal and Montani, 2023) and the TensorFlow v.2.12.0 (Abadi et al., 2016) spaCy Universal Sentence Encoder v.0.4.6 (Mensio, 2023) were used to process the transcripts to obtain semantic similarity scores. Three phrases, “tall”, “tall plant”, and “tall height”, were compared to each row observation through spaCy’s similarity function using the pre-trained large English universal sentence encoder (en use lg) from TensorFlow. A dataset of similarity scores in the form of values from 0 to 1 was generated (Figure 2b). The spaCy Universal sentence encoder and the similarity function we used scores semantic similarity from 0 to 1 based on the phrase used to compare the observations. Therefore, we selected to use all tall terms as the query (rather than also focusing on phrases for short stature) so that observations for tall plants would be closer to a score of 1 (observations meaning the opposite, i.e. small plants would be scored closer to 0). If the word short were substituted for tall as the query, then smaller would be scored closer to 1 and tall plants closer to 0.

Similarity scores for the 653 taxa shared by both datasets were retained, encompassing 35,709 rows or 91.92% of the original rows observed (Table 2). The similarity scores for the “tall” query were used as input to calculate BLUEs and BLUPs in the same manner as described in the *Plant Height Measurement Data* section, and visualizations of diagnostics plots of the models (Supplementary Figure 3, Supplementary Figure 4) were generated.

**Table 2.**
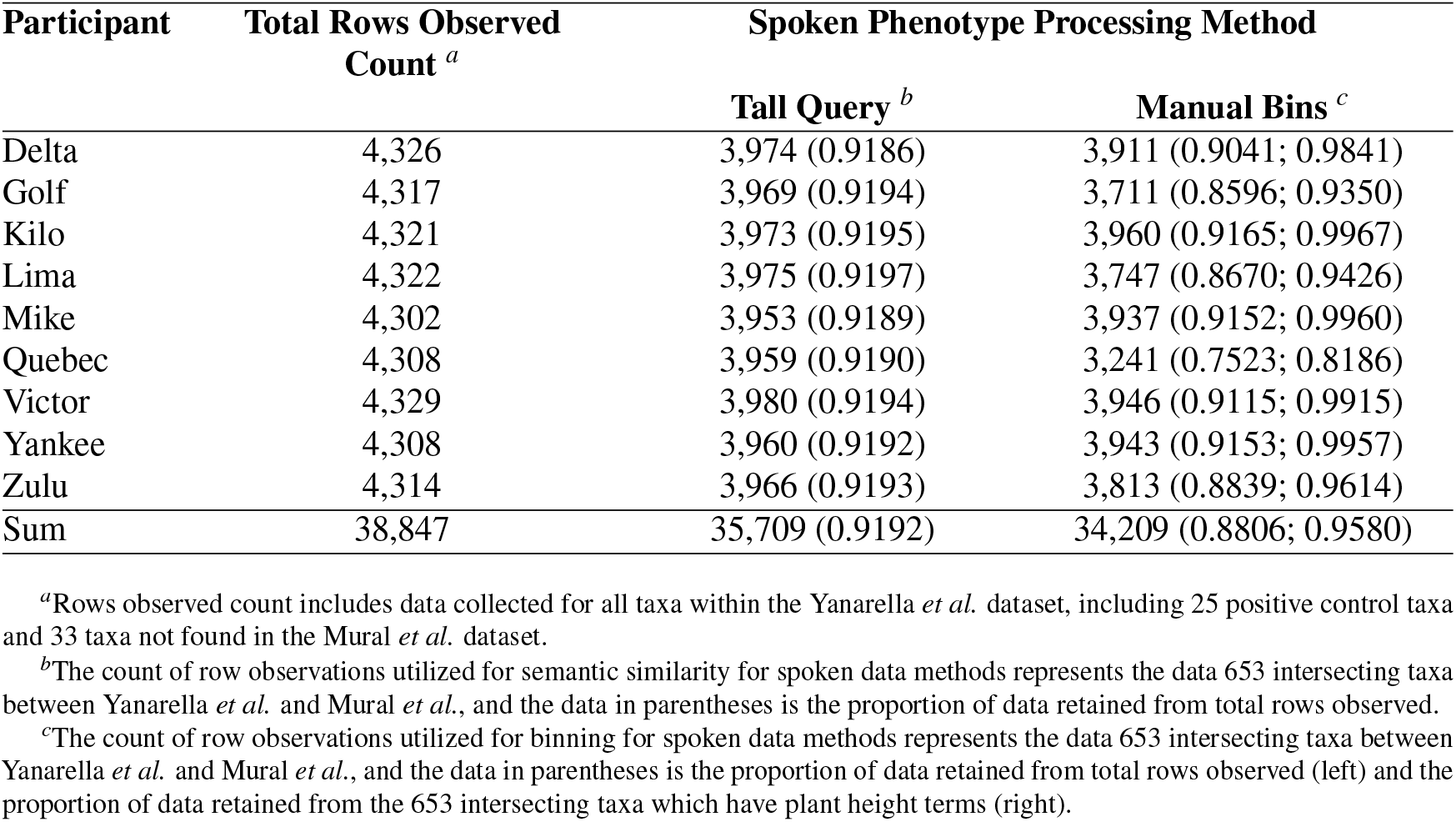
Observation retention by spoken phenotype method.

| Participant | Total Rows Observed<br>Count <sup>a</sup> | Spoken Phenotype Processing Method |  |
| --- | --- | --- | --- |
|  |  | Tall Query <sup>b</sup> | Manual Bins <sup>c</sup> |
| Delta | 4,326 | 3,974 (0.9186) | 3,911 (0.9041; 0.9841) |
| Golf | 4,317 | 3,969 (0.9194) | 3,711 (0.8596; 0.9350) |
| Kilo | 4,321 | 3,973 (0.9195) | 3,960 (0.9165; 0.9967) |
| Lima | 4,322 | 3,975 (0.9197) | 3,747 (0.8670; 0.9426) |
| Mike | 4,302 | 3,953 (0.9189) | 3,937 (0.9152; 0.9960) |
| Quebec | 4,308 | 3,959 (0.9190) | 3,241 (0.7523; 0.8186) |
| Victor | 4,329 | 3,980 (0.9194) | 3,946 (0.9115; 0.9915) |
| Yankee | 4,308 | 3,960 (0.9192) | 3,943 (0.9153; 0.9957) |
| Zulu | 4,314 | 3,966 (0.9193) | 3,813 (0.8839; 0.9614) |
| Sum | 38,847 | 35,709 (0.9192) | 34,209 (0.8806; 0.9580) |

<sup>a</sup>Rows observed count includes data collected for all taxa within the Yanarella *et al.* dataset, including 25 positive control taxa and 33 taxa not found in the Mural *et al.* dataset.
<sup>b</sup>The count of row observations utilized for semantic similarity for spoken data methods represents the data 653 intersecting taxa between Yanarella *et al.* and Mural *et al.*, and the data in parentheses is the proportion of data retained from total rows observed.
<sup>c</sup>The count of row observations utilized for binning for spoken data methods represents the data 653 intersecting taxa between Yanarella *et al.* and Mural *et al.*, and the data in parentheses is the proportion of data retained from total rows observed (left) and the proportion of data retained from the 653 intersecting taxa which have plant height terms (right).

#### 2.3.3 Binning for Plant Height Spoken Data

The transcripts of spoken plant descriptions were mined manually and programmatically for phrases directly related to narrations about plant height. A set of 797 plant height phrases were identified and manually curated and binned from 0 to 7, where 0: no growth, 1: very short plants, 2: short plants, 3: short-medium height plants, 4: medium height plants, 5: medium-tall height plants, 6: tall plants, 7: very tall plants. Bin values were assigned to observations for the 653 taxa shared by both datasets (Figure 2b). Of the total text transcripts, there were 34,209 or 88.06% row observations that were retained and binned (Table 2).

BLUEs and BLUPs were calculated as described in the *Plant Height Measurement Data* Section, and visualizations of diagnostics plots of the models (Supplementary Figure 5, Supplementary Figure 6) were generated. Additionally, the R nnet v.7.3-19 package (Venables and Ripley, 2002) was used to perform multinomial logistic regression to predict height phrase bins, where taxa and replicate were fit as fixed effects.

### 2.4 Preprocessing the Genotypic Dataset

To ease the subsetting step to obtain the data for the 653 that overlap between the Mural and Yanarella datasets, Trait Analysis by aSSociation, Evolution and Linkage (TASSEL) Version 5.0 Standalone (Bradbury et al., 2007) was used to convert the Mural *et al*. genotypic data from a variant call format (vcf) formatted file to a HapMap (hmp) formatted file. The data was then processed to contain marker information for the 653 taxa shared with the Yanarella *et al*. dataset, these data were grouped by chromosome (because datasets were so large as to exceed the limits of the HPC memory we were able to utilize), and HapMap files were generated for each chromosome. The chromosome files were sorted by maker position from lowest position to highest position.

The sorted chromosome files were reformatted to vcf files using TASSEL Version 5.0 Standalone, then vcftools v.0.1.14 (Danecek et al., 2011) concatenated these files, and the resulting file was zipped. The concatenated data (a subset of 653 lines) were then again converted to a VCF format for use with PopLDdecay v3.42 (Zhang et al., 2018) to analyze and visualize Linkage Disequilibrium (LD) Decay of the Mural *et al*. genotypic data.

### 2.5 Genome-Wide Association Studies and Analysis

Genome Association and Prediction Integrated Tool (GAPIT) 3 v.3.1, 2022.4.16 (Lipka et al., 2012; Tang et al., 2016; Wang and Zhang, 2021) was used to perform Fixed and random model Circulating Probability Unification (FarmCPU) (Liu et al., 2016) and Mixed Linear Model (MLM) (Yu et al., 2005) on each of the phenotypic datasets using the Mural *et al*. marker dataset for genotypic input. Each chromosome was run individually, and PCA.total parameter was set to 3 for all analyses (Figure 2c). This manuscript focuses on the FarmCPU analyses, the MLM processing and results are available as described in *Web resources* section.

Manhattan plots from the resulting GAPIT analyses were generated using the ggplot2 v.3.4.3 package (Wickham, 2016). We used the RAINBOWR v.0.1.29 package’s (Hamazaki and Iwata, 2020) CalcThreshold to determine the Bonferroni threshold for each analysis with a sig.level of 0.05. SNPs that were identified as above the Bonferroni threshold for each analysis were viewed on MaizeGDB’s implementation of GBrowse2 (Generic Genome Browser v.2.55) (Stein, 2013) using Maize B73 RefGen v4; gene IDs were collected within +/-300 kilobases (kb), based on the LD decay curve generated by PopLDdecay, of the identified SNPs (Supplementary Figure 7).

We collected a list of genes shown to influence plant height (Table 3, Supplementary Table 2) and compared the gene IDs from the GWAS analyses using a web-based intersection and Venn diagram tool (Sterck, 2021) to determine if these previously published plant height genes were identified within the +/-300 kb region indicated by the LD decay curve for each of our analyses. Additionally, Gene Ontology (GO) terms for the gene IDs within +/-300 kb of SNPs identified as significant were obtained using the B73 RefGen V4 Zm00001d.2 annotations generated by Maize GO Annotation -Methods, Evaluation, and Review (maize-GAMER) tool (Wimalanathan and Lawrence-Dill, 2017; Wimalanathan et al., 2018) and the R package GO.db v.3.17.0 (Carlson, 2023) was used to collect terms associated to the GO IDs (Supplementary Table 3). Where no GO terms were associated with genes or gene models or those associated were not clearly related to the plant height trait, knowledge of gene function via associated *Arabidopsis thaliana* orthologues listed in MaizeGDB and linked to TAIR were reviewed for possible insight (Andorf et al., 2016; Reiser et al., 2022).

**Table 3.** Plant height gene models count identified from publications.

| Publication | Plant Height Related Gene Model Count<sup>a</sup> |
| --- | --- |
| Jansson (1994) | 1 |
| Winkler and Helentjaris (1995) | 1 |
| Austin and Lee (1996) | 3 |
| Multani et al. (2003) | 2 |
| Blakeslee et al. (2005) | 1 |
| Geisler and Murphy (2005) | 1 |
| Brooks et al. (2009) | 1 |
| Wu et al. (2009) | 7 |
| Bai et al. (2009) | 6 |
| Lawit et al. (2010) | 2 |
| Hartwig et al. (2011) | 1 |
| Salvi et al. (2011) | 4 |
| Weng et al. (2011) | 1 |
| Teng et al. (2012) | 1 |
| Gallavotti (2013) | 1 |
| Peiffer et al. (2014) | 7 |
| Wallace et al. (2016) | 312 |
| Mazaheri et al. (2019) | 10 |
| Azodi et al. (2020) | 876 |
| Mural et al. (2022b) | 347 |

<sup>a</sup>Count of gene model IDs identified for Zm00001d.2 gene model and Zm-B73-REFERENCE-GRAMENE-4.0 assembly version.

## 3 RESULTS AND DISCUSSION

Student participants made three complete passes of the field (∼4,320 observations per student participant (Table 2, *Total Rows Observed Count*) and used their individual wording and phraseology to describe phenotypes. Recorded observations varied from 2 to 241 words in length (Figure 3). From this dataset (described fully in Yanarella et al. et al., 2024), all analyses and results reported here are derived.

**Figure 3.**
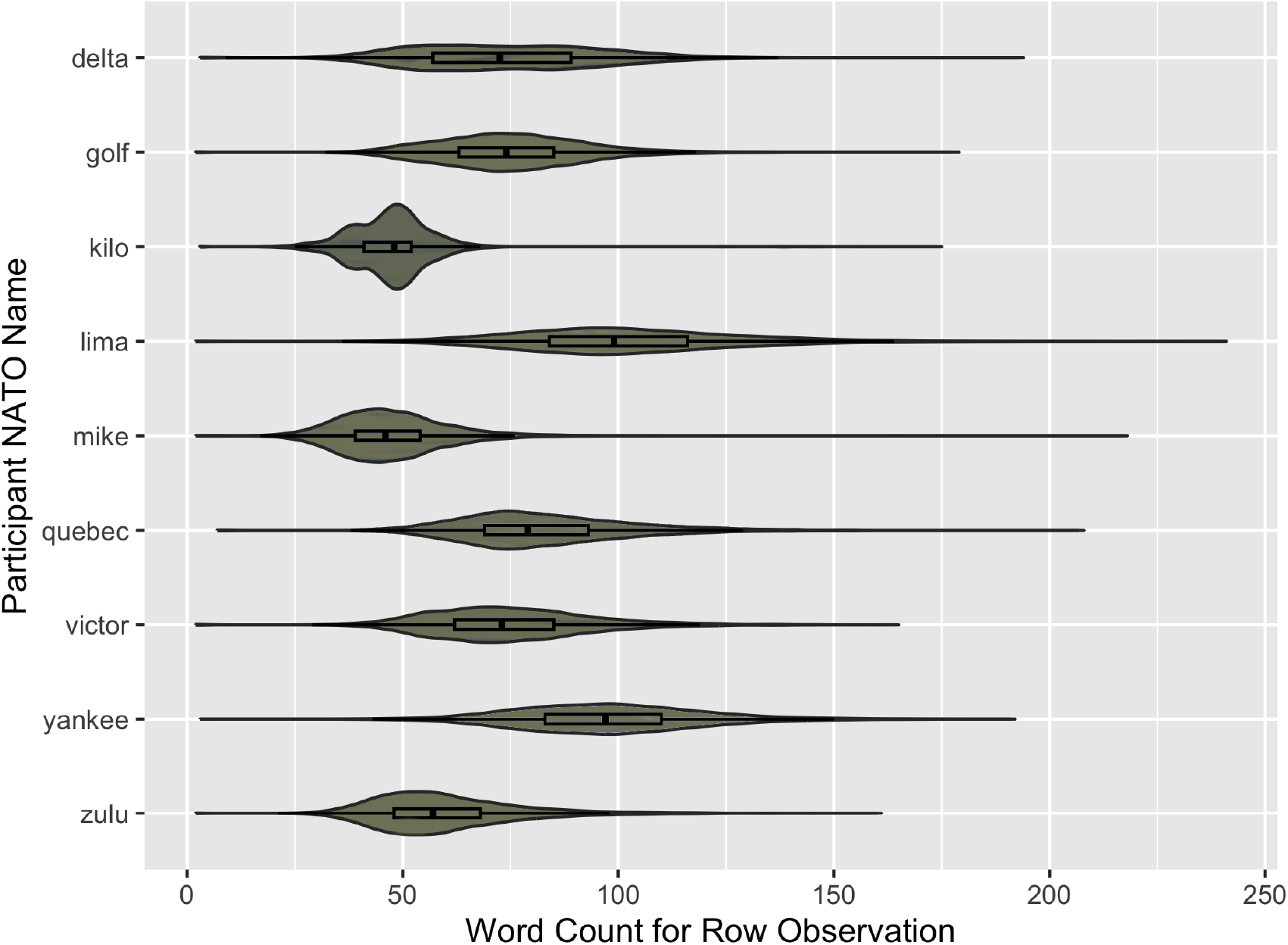
Distributions of each student participant’s word count per observation. X axis shows number of words for individual row obeservation. Y axis indicates each participant’s NATO name. Boxplots are included within each of the violin plots, individual outlier points not represented.

### 3.1 Visually Striking Phenotypes are Detected from Unstructured Spoken Descriptions

We reasoned generally that including visually observable phenotypes with known phenotype-gene associations would be a first step toward understanding whether individuals without specific training on describing plant phenotypes could recover known phenotype-gene associations. To explore student participants’ ability to identify and describe phenotypes, 25 positive control accessions that, if grown in the appropriate environmental conditions, would show visually “dramatic” color phenotypes were included in the field.

The 25 positive control accessions were observed 53-55 times over all nine student participants (Table 1). We utilized Merriam-Webster (2023) and WordHippo (Kat IP Pty Ltd, 2008) thesaurus services to determine participants’ ability to identify words synonymous with descriptors that characterize positive control phenotypes as demonstated in Figure 4a-c. For example, accessions M241C *A1 A2 B1 C1 C2 Pl1 Pr1 R1-r* and 219L *B1-S; R1-r pl1-McClintock* (gene name *colored1* and *colored plant1*, Supplementary Table 1) had at least one synonym in each of the observations made by the student participants as indicated in Table 1 by the proportion of 1.000 for both Merriam-Webster and WordHippo synonyms. While accessions U740G *Fbr1-N1602* (gene name *few branched1*), 703J *Rs1-O 1* and 703K *Rs1-Z* (gene name *rough sheath1*, Supplementary Table 1) had low proportions of observations having at least one synonym for each observation as indicated in Table 1.

**Figure 4.**
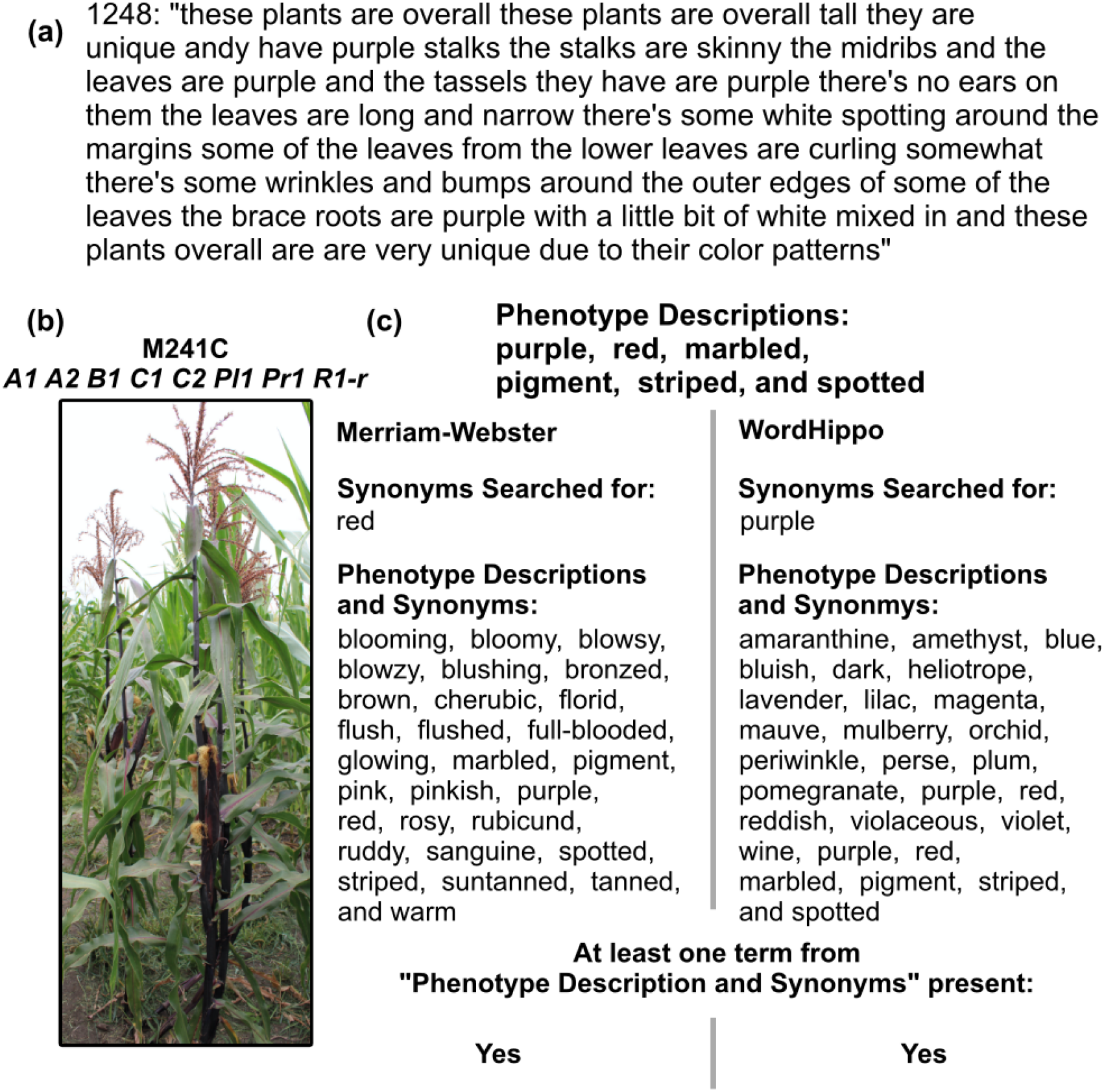
Example of detecting phenotypes from positive control accession descriptions. (a) Transcript of a spoken description for positive control accession M241C *A1 A2 B1 C1 C2 PI1 Pr1 R1-r*, (b) Image of positive control accession M241C *A1 A2 B1 C1 C2 PI1 Pr1 R1-r*, (c) Example of phenotype description terms, and synonyms from Merriam-Webster and WordHippo, to demonstrate a description having at least one instance of a synonymous word for the phenotype of interest.

Our findings indicate that non-expert participants, unaware of expected phenotypes, can identify and describe what they observe using terms that could be valuable for genomic association studies. These findings are, however, constrained by the number of synonyms used for phenotype descriptions and the specific environmental conditions of the plant growth. The reliance on thesaurus services could also potentially reduce the proportion of observations with at least one synonym, particularly if participants opted for informal descriptions. Furthermore, accurate descriptions of our intended positive control phenotypes were contingent upon favorable field environment and weather conditions.

### 3.2 Plant Height Assessments can be Derived from Unstructured Spoken Descriptions

Given that visually apparent phenotypes were described successfully by non-expert participants for control lines grown in the field, we moved on to assessing whether plant height, a continuous and subjective trait from ‘short’ to ‘tall’, could be derived from the WiDiv population, a large association mapping panel.

Two methods were employed to pre-process text transcriptions of spoken descriptions of plants. The semantic similarity method of comparing the term “tall” to each row observation retained 91.92% of the full set of row recordings captured (including the 25 positive controls and 33 accessions unique to the Yanarella *et al*. dataset) by the student participants, demonstrating that 35,709 row observations were made with taxa in both datasets (Table 2). The manual bin method of identifying phrases related to plant height and binning them based on apparent semantic similarity retained 86.06% of the full set of row recordings captured by the student participants and 95.80% of the observations with plant height phrases made with taxa in both datasets, which results in 34,209 row observations for manually binned data (Table 2).

Both methods parse information about the plant height trait and process the data into a format appropriate as input into available GWAS tools and models. Using a query term for plant height and semantic similarity requires less manual curation and was implemented on a larger subset of data. The benefit of the binning method is that it reduces the noise (where noise in the context of natural language processing refers to additional information not directly related to the particular topic of interest). Only observations with plant height-related terms were considered.

A limitation of the query term and semantic similarity method is retaining noisy data because this method compares the “tall” query to each observation and relies on pre-trained models. An example of noise comes from the participant NATO code name “Victor”. Their recording for row 1,456 on 07/16/2021, *tall and height green all the way to the bottom* … *super short hairs on top that are quite prickly* … *in general there are the brace roots are short fat and light green in color*, in which the similarity function of spaCy University Sentence Encoder (Mensio, 2023) when compared to the “tall” query string determine the semantic similarity score as 0.0848. The shortcomings of the binning method are the time-consuming nature of curating lists of phrases relevant to the trait of interest and the loss of data where phrases directly related to plant height were not specified.

### 3.3 Genomic Associations for Plant Height can be Derived from Unstructured Spoken Descriptions of Phenotype

Efforts to assess whether spoken data could be used for GWAS were structured as follows. We collected a dataset of manually measured plant heights, which would serve as ground truth for assessing any language-based findings. We also processed spoken descriptions of lines in two ways (semantic similarity versus a binning method), and carried out GWAS using BLUEs and BLUPs to consider differences in random and fixed effects across the variables taxon, replicate, and row number. Once regions of the genome associated with the trait plant height were derived for manually measured, semantic similarity, and binned datasets, significant regions that included genes known to be involved in plant height were highlighted. Loci identified as significant from both spoken datasets were compared to the manual, ground truth dataset to determine whether the same regions were identified across both data collection methods, and all genes in regions of significant association for the plant height trait were assessed for GO terms associated with plant growth hormones auxin, brassinosteroid, and gibberellin.

#### 3.3.1 Ground Truth: Manual Plant Height Measurement GWAS

We performed association studies using FarmCPU (Liu et al., 2016) on three categories of phenotype data. The first phenotype category was ground truth (measured) plant height data using BLUEs (Figure 5a) and BLUPs (Figure 5b). For the BLUE analysis, 21 significant SNPs above the Bonferroni threshold of 8.55 were identified, and of those SNPs, we discovered 10 (Supplementary Table 3; colored orange) in which at least one plant height gene was detected in the literature within +/-300 kb (Supplementary Table 2). The BLUP analysis identified 29 significant SNPs, 9 (Supplementary Table 3) where at least one plant height gene was discovered in the literature within +/-300 kb (Table 3, Supplementary Table 2).

**Figure 5.**
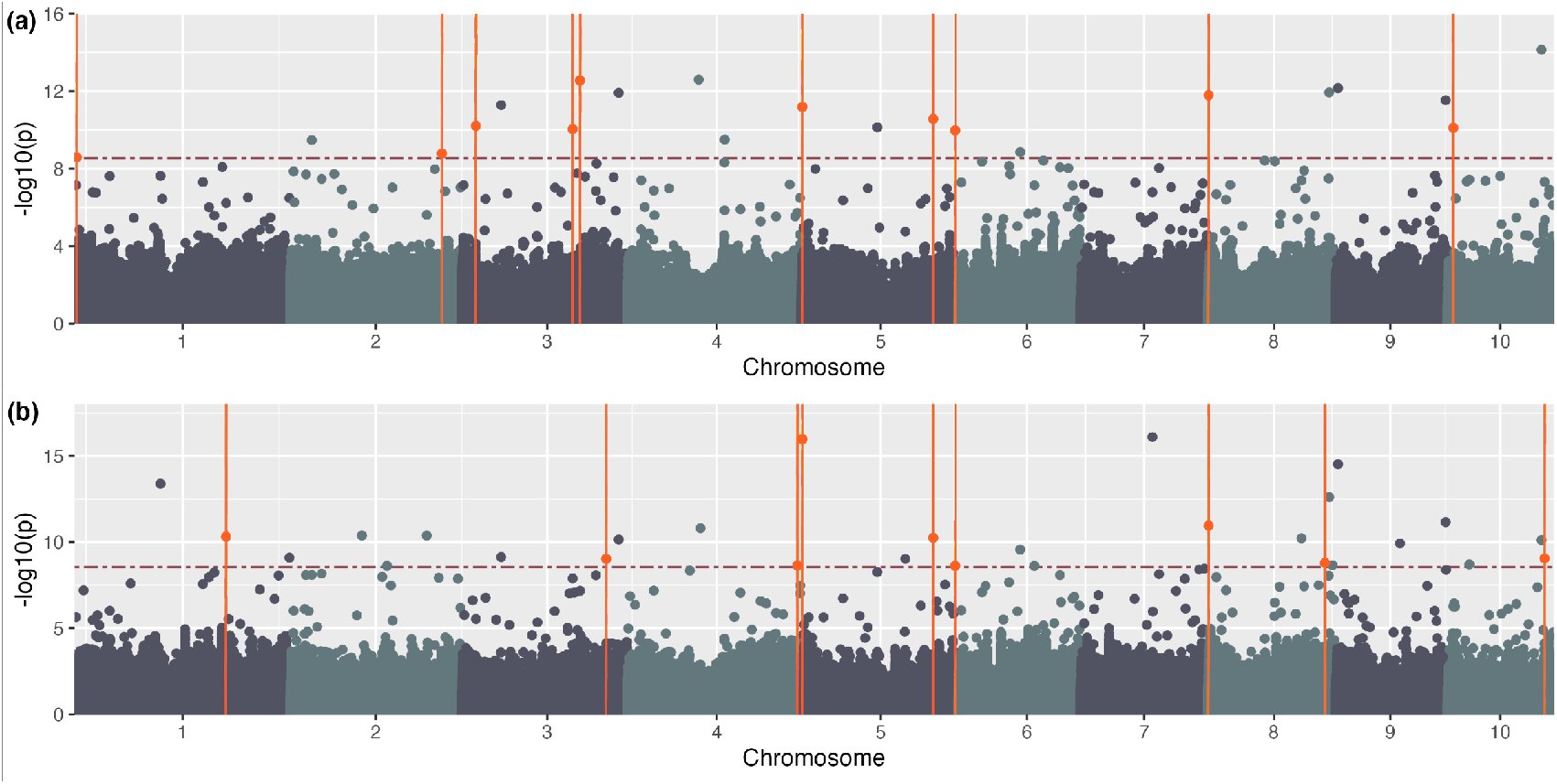
Manually measured height -phenotypic data. Manhattan plot generated using GAPIT and FarmCPU using measured height data (a) BLUEs and (b) BLUPs using Mural *et al*. genotypic data. The horizontal red dashed line indicates the Bonferroni threshold ((a) -log 10(p) = 8.55; (b) -log 10(p) = 8.55); orange points above the line and intersected by a vertical line indicate identified SNPs with known plant height genes within +/-300 kb.

#### 3.3.2 Semantic Similarity for Plant Height Spoken Data GWAS

The second phenotype category used BLUEs (Figure 6a) and BLUPs (Figure 6b) for tall query and semantic similarity of spoken phenotype descriptions. These analyses identified 27 and 23 significant SNPs (Supplementary Table 3; colored orange), respectively, above the Bonferroni threshold of 8.55. Of these, 9 and 8 genes, respectively, were formerly detected for plant height within +/-300 kb of the SNP (Table 3, Supplementary Table 2).

**Figure 6.**
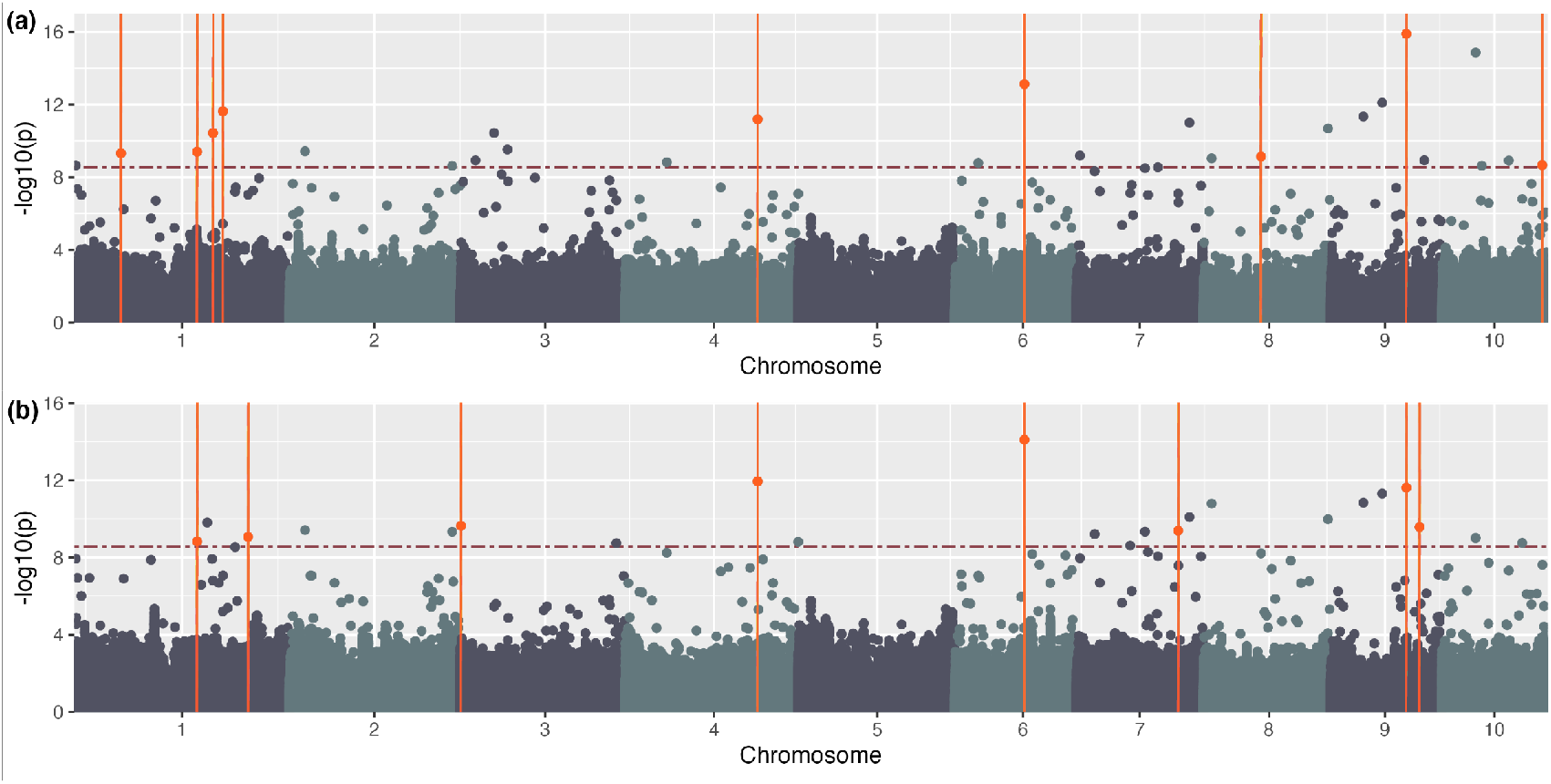
Semantic similarity for query tall -phenotypic data. Manhattan plot generated using GAPIT and FarmCPU using semantic similarity score data (a) BLUEs and (b) BLUPs using Mural *et al*. genotypic data. The horizontal red dashed line indicates the Bonferroni threshold ((a) -log 10(p) = 8.55; (b) -log 10(p) = 8.55); orange points above the line and intersected by a vertical line indicate identified SNPs with known plant height genes within +/-300 kb.

The semantic similarity method with the tall query to generate phenotype values for each spoken observation using spaCy’s similarity function (Honnibal and Montani, 2023) and the Universal Sentence Encoder (Mensio, 2023) was successful. We were able to perform GWAS, and there were significant SNPs from regions associated with plant height. As this is a proof of concept study, we acknowledge that other pre-trained models exist capable of calculating semantic similarity or models that can be adapted to generate similarity scores related to plant height such as BioBERT (Lee et al., 2019), those implemented by the Python gensim package (Řehůřek and Sojika, 2010), or others reviewed in Koroleva et al. (2019). Further, additional queries could be employed for relating the text observations to a height value.

#### 3.3.3 Binning for Plant Height Spoken Data GWAS

The third phenotype category used BLUEs (Figure 7a) and BLUPs (Figure 7b) for manual binning of spoken phenotype descriptions with plant height terms. These analyses identified 32 and 33, respectively, significant SNPs (Supplementary Table 3; colored orange), above the Bonferroni threshold of 8.55. Of these, 13 and 12 have genes formerly reported within +/-300 kb of the SNP (Table 3, Supplementary Table 2). An additional analysis was completed using predicted values from a multinomial regression (Supplementary Figure 8, Supplementary Table 2) in which 21 SNPs were significant, and 3 had genes detected within +/-300 kb of the SNP in the literature (Table 3, Supplementary Table 2).

**Figure 7.**
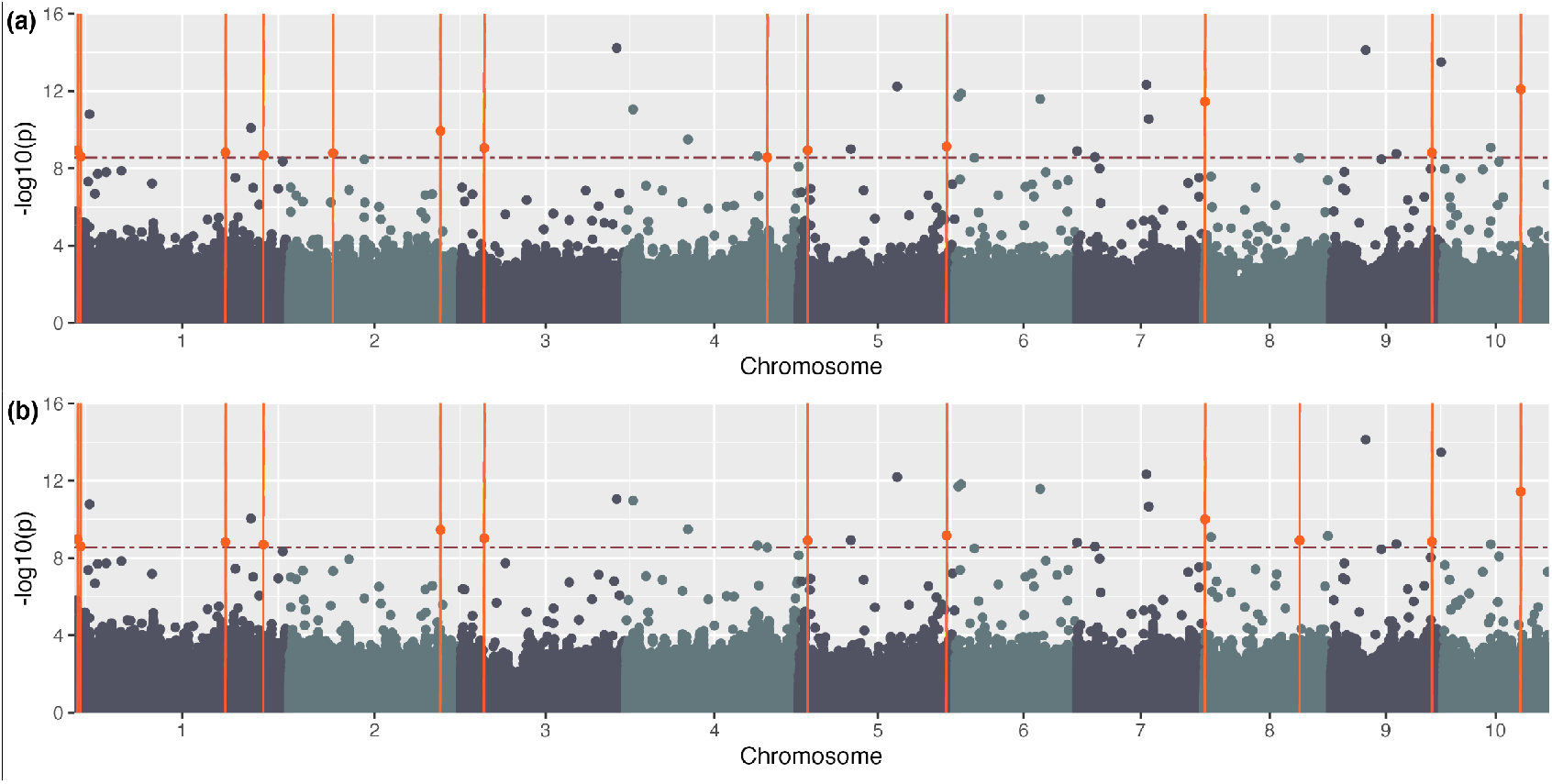
Binned phrases for plant height - phenotypic data. Manhattan plot generated using GAPIT and FarmCPU using binned plant height data (a) BLUEs and (b) BLUPs using Mural *et al*. genotypic data. The horizontal red dashed line indicates the Bonferroni threshold ((a) -log 10(p) = 8.55; (b) -log 10(p) = 8.55); orange points above the line and intersected by a vertical line indicate identified SNPs with known plant height genes within +/-300 kb.

The binning method for plant height phrases appears to be a promising method for association studies with phenotype data extracted from spoken descriptions. This method reduces the noisiness of the transcription data and scores observations on only phrases detailing features of plant height. Additionally, the GWAS performed with BLUE and BLUP values generated from the binning method detected more known regions associated with plant height formerly reported than the manually measured and semantic similarity query methods.

#### 3.3.4 Comparing Across Manual Measurement, Semantic Similarity, and Binning

Within the +/-300 kb region indicated by the LD decay curve for each of the analyses, we lined up shared significant regions within BLUEs and BLUPs for each of the three studies (i.e., manual measurement, semantic similarity, and binning). These comparisons are shown in Supplementary Table 4. Within the manual measurements, 11 loci identified are shared across BLUEs and BLUPs. For semantic similarity, 12 shared regions are identified. For binning, 30 shared regions are identified.

Of particular interest are comparisons between manual measurement and the two methods derived from unstructured, spoken language descriptions of phenotype (i.e., semantic similarity and binning). Manual measurement and semantic similarity analyses showed no correspondence. However, manual measurement and binning shared four regions for BLUEs, with one of the four regions resulting from the same SNP (as opposed to the other three, where different SNP within the +/-300 kb region are responsible for the correspondence). Four SNPs in that shared group (representing a region on Chromosome 1 and a region on Chromosome 8) contain genes known to be associated with plant height from the literature (marked in orange in Supplementary Table 4). For BLUPs, manual measurement and binning share six regions, with three regions resulting from shared SNP (and three as a result of different SNP within the +/-300 kb region). Four SNPs in that shared group (representing a region on Chromosome 1 and a region on Chromosome 8) contain genes known to be associated with plant height from the literature (likewise indicated in orange). The significant shared SNPs on chromosome 8 for each group are shared across measured data binned data for both BLUEs and BLUPs.

Of some note are the regions that are *not* shared between manual measurements and the semantic similarity or binning GWAS outputs. For those significant associations that are only present in the semantic similarity and/or binning outputs, some of these candidate genes for plant height agree with other studies. For others, they are not documented in other studies, making them truly novel candidate genes identified via association genetics based on spoken phenotypic descriptions.

#### 3.3.5 Example Candidate Genes in Select Regions

To give examples of how to review and prioritize candidate genes for the plant height trait in some of the regions identified, we describe here the three regions that were shared between the measured and binned results sets for both BLUEs and BLUPs. These constitute three loci, one each on chromosomes 3, 8, and 9. Shown in Table 5 are these three loci, of which one (on chromosome 8) coincides with a previously published association from GWAS for plant height (red), and the other two are newly identified through GWAS in this study from spoken phenotypic data collection techniques as well as from the manual scoring procedure.

Table 4 shows the SNPs identified, the number of gene models in the +/-300 bp region that contains the SNPs. Where characterized genes are known for those gene models, gene model names as well as gene names and symbols are also listed. GO terms relevant to the plant height trait are shown in bold. For the chromosome 3 region, no gene models in the region had any GO terms obviously related to height, so we also looked up the gene models via MaizeGDB to find out whether any associated *Arabidopsis* orthologues might have known phenotypes or functions related to plant height.

**Table 4.**
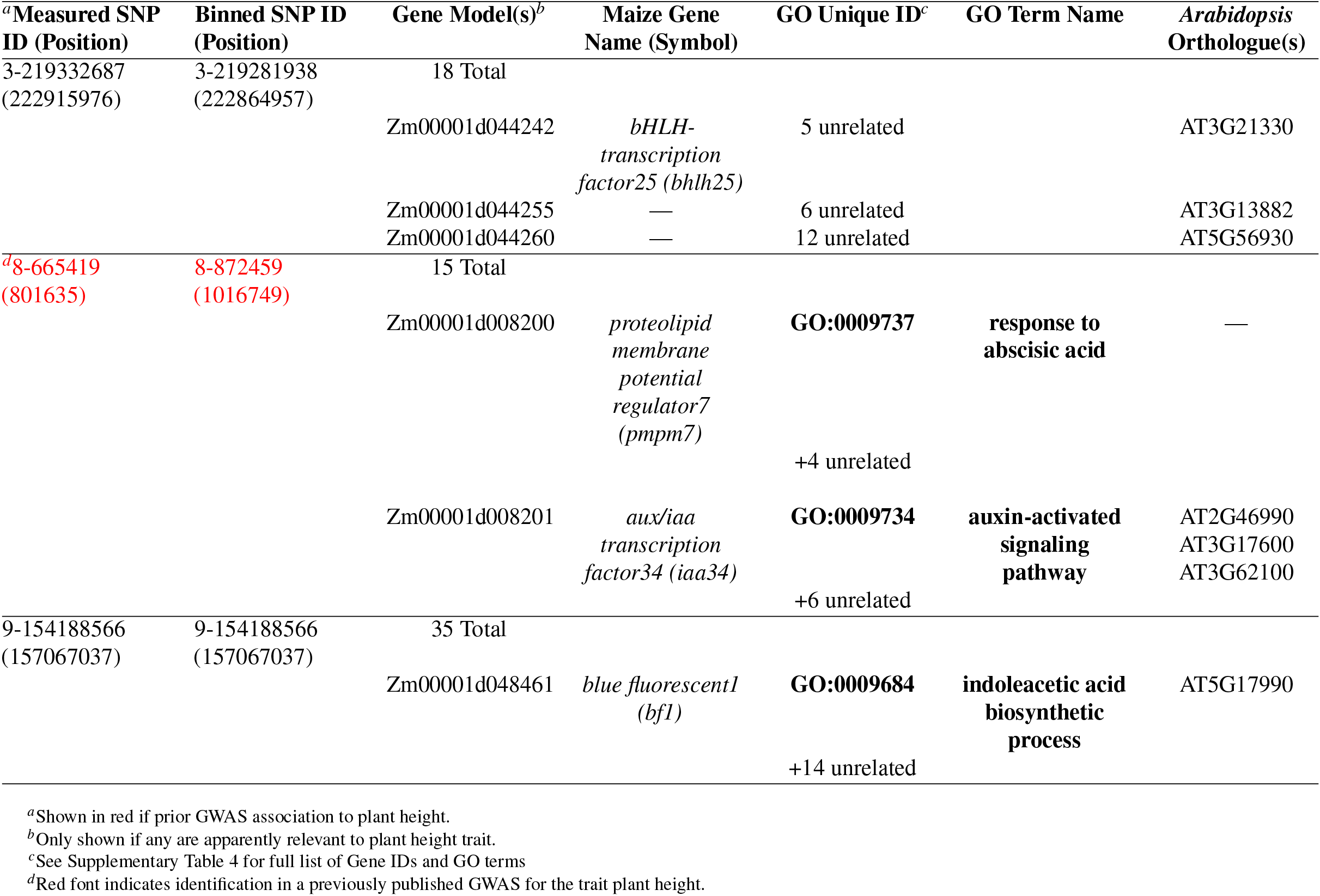
Candidate genes for plant height, shared measured and binned BLUEs and BLUPs.

Promising candidate genes are as follows. For the newly identified chromosome 3 locus, three candidate gene models are: Zm00001d044242, the gene *bhlh25*, which has been implicated in abscisic acid biosynthesis/regulation (Vendramin et al. et al. 2020; and *Arabidopsis* orthologue AT3G21330 is a member of a family of genes involved in photoresponsiveness for hypocotyl length; Khanna et al., et al. 2006), Zm00001d044255 (*Arabidopsis* orthologue AT3G13882 is involved in plant growth rates and flowering; Xu et al., 2023), and Zm00001d044260, the gene *c3h2*, with *c3h* genes showing some height-related phenotypic involvement (Fornalé et al., et al. 2015; and *Arabidopsis* orthologue AT5G56930, also called AtC3H65, is involved in brassinosteroid signaling; Wang et al., 2022). For the chromosome 8 locus previously associated with plant height by Azodi et al. (2020), the loci *pmpm7* (also called PMP3-7 in some literature) and *iaa34* are the only genes in the region. Both are known to be responsible for plant growth and height phenotypes (Fu et al., 2012; Galli et al., 2015) and are associated with GO terms that are likely relevant. For the newly identified chromosome 9 locus, one candidate gene model looks promising: Zm00001d048461 with associated term GO:0009684, indoleacetic acid biosynthetic process (where indoleacetic acid or IAA is an auxin). Upon further investigation, this gene model represents the gene *blue fluorescence1* (*bf1*) known to affect maize plant stature (reviewed by Khavkin and Coe, et al. 1997), but not among the literature of associated plant height genes we had assembled for this work. Note: for all loci identified as being significantly associated with the trait plant height, the information that would enable these same analysis steps can be accessed via Supplementary Tables 3 and 5.

#### 3.3.6 Plant Growth Hormone Functions in Genomic Regions Associated with Plant Height

To assess relationships to plant growth hormones and identified gene IDs across the whole genome, we reviewed the set of GO terms annotated to gene IDs associated with the regions +/-300 kb of significant SNPs. The full list of GO terms for each model is available in Supplementary Table 3 and Supplementary Table 5. To examine how these terms align with plant height terms, we queried the dataset for the words auxin, brassinosteroid, and gibberellin because of their known functions in plant height regulation (Li et al., 2020 reviews the importance of these hormones). The term “auxin” was more frequently present in these datasets when compared to brassinosteroid or gibberellin (Table 5).

**Table 5.** Appearance of GO terms by plant hormone for each method.

| Analysis | auxin |  | brassinosteroid |  | gibberellin |  |
| --- | --- | --- | --- | --- | --- | --- |
|  | Literature | <sup>a</sup> This Study | Literature | This Study | Literature | This Study |
| Measured BLUEs GO Terms | 2 | 15 | 0 | 3 | 0 | 4 |
| Measured BLUPs GO Terms | 3 | 22 | 0 | 8 | 0 | 6 |
| Semantic Similarity Tall Query BLUEs GO Terms | 12 | 5 | 0 | 5 | 0 | 7 |
| Semantic Similarity Tall Query BLUPs GO Terms | 2 | 15 | 0 | 3 | 0 | 6 |
| Manual Term Bin BLUEs GO Terms | 1 | 16 | 0 | 8 | 1 | 10 |
| Manual Term Bin BLUPs GO Terms | 1 | 15 | 0 | 6 | 1 | 9 |
| Manual Term Bin Multinomial GO Terms | 0 | 10 | 0 | 3 | 0 | 1 |

<sup>a</sup> Appearance count for GO terms of prior literature in Table 3.
<sup>b</sup> Appearance count for GO terms unique to this study.

Additionally, other GO annotations identified in our analyses have functions that may affect plant height. Examples of these GO terms include developmental growth (GO:0048589), anatomical structure formation involved in morphogenesis (GO:0048646), and shoot system development (GO:0048367). Further, using GWAS, we found genomic regions with predicted functional annotations related to plant hormone functions that were not reported in the literature in Table 3. These regions may be of interest for follow-on experimentation to assess potential involvement in the plant height trait.

While examining GO terms, descriptions that do not relate to plant functions but were annotated to gene IDs occurred. Errant assignments of GO terms for plant-specific tasks has been described in Fattel et al. (2022). An example is the gene ID *Zm00001d008201* being assigned the term animal organ development (GO:0048513). Interestingly, this ID was also assigned the terms auxin-activated signaling pathway (GO:0009734) and post-embryonic development (GO:0009791). These results demonstrate a compelling argument for reviewing GO terms carefully.

#### 3.3.7 Additional Methods Tried and Room for Improvement

We demonstrate the use of a multinomial regression to generate phenotypic input with binned data, although we recognize that FarmCPU is not optimized for multinomial input. GWAS tools that utilize an ordered multinomial regression model to predict multinomial values for association studies were developed in the medical research field (German et al., 2019). Regardless, FarmCPU with multinomial binned input for plant height detected regions of the genome associated with plant height.

While participant language was not constrained, input that is less noisy and with lower data loss could be attainable if an emphasis were placed on stating specific aspects of the plant accompanied by a descriptor and reducing literary descriptive comments. An example of describing a specific aspect of a plant is “tall, green, and long” compared to “this row has tall plants.” The former statement is unclear whether the whole plant is described or a specific aspect of the plant, while the latter makes it clearer that the total plant height is described. Literary language descriptions are more difficult to compute because context is necessary to determine the meaning behind a phrase, an example, candy cane stripe. While candy cane stripe may induce a mental image of a candy cane, unless a computation model is trained to identify the literary description, the model would not be able to discern the spoken description of phenotype as a particular striped pattern.

## 4 CONCLUSION

By translating the linguistic complexity of human speech into data compatible with GWAS tools, we have demonstrated a successful proof of concept that paves the way for further innovation in data collection and analysis. Our two-fold approach—semantic similarity and manual binning—proved to be robust, capturing known genetic associations with plant height and identifying new regions of interest. Of note, those describing the plants were not aware that plant height would be a focus of the study, and no guidance on what was “tall” or “short” was provided. Nonetheless, known genetic associations with the trait plant height were uncovered. These findings underscore the potential for expanding the scope of phenotypic data collection methods in genetic studies. By confirming that unstructured spoken data can yield quantifiable results for genetic association, we open the door to complementary and diverse data collection techniques that may be more tractable for non-expert involvement. This could enable the inclusion of non-experts in data collection, and significantly enrich the dataset by bringing in previously overlooked phenotypic nuances.

Beyond these demonstrations that spoken, unstructured phenotypic descriptions can be used to recover known associations, there are two other conceptual benefits that should be considered. Firstly, when people are describing what they see in the field rather than exclusively collecting predefined traits, the potential to uncover novel phenomena is perhaps increased. Secondly, it is the case that for many years we have used computers to analyzed structured data, so those collecting the data have limited themselves to documenting data in a structured, computer-friendly format. This is, in effect, asking people to structure their thinking and documentation like - and for - a computer. With the methods described here, the people collecting the data are enabled to think and behave in a more naturally human way for data collection. This has implications for the rate of data collection and for cognitive burden as follows. Over three weeks of data collection, each participant made three complete passes of the field, recording spoken observations. However, for manual scoring and measurement data, none were able to make a single complete pass of the field. Our experimental design enabled the student participants to speak and describe plant traits using their unique vocabulary and speech patterns. Participant “Zulu” reported that recording spoken observations was simpler and easier than measuring and scoring because they could make more detailed observations about different parts of the plants because recording spoken observations was both less strenuous and less mentally taxing, indicating that data collection through speech may reduce the cognitive load on field researchers.

While image-based data collection derived from unmanned vehicles equipped with sensors for visual phenotype detection continues to improve and is invaluable for certain traits (reviewed in Xiao et al., 2022), it falls short for traits requiring, e.g., tactile or olfactory observations. Coupling spoken and written language-based annotations with image analysis can enhance our understanding of complex phenotypes, harnessing human perception to capture nuanced details that purely image-based approaches might miss. This work demonstrates that such data can be collected in straightforward ways, and that phenotypic information is indeed accessible for large-scale genetics and genomics analyses.

## 5 WEB RESOURCES

Code to recreate the analysis in this manuscript is available at CyVerse Data Commons from (Yanarella et al., 2023b) and can be accessed from: https://datacommons.cyverse.org/browse/iplant/home/shared/commons_repo/curated/Carolyn_Lawrence_Dill_Maize_WiDiv_Association_Studies_Dataset_September_2023.

## 6 DATA AVAILABILITY

The de-identified spoken data described in this manuscript is exempted by Iowa State University’s Institutional Review Board (IRB ID: 21-179-00). Phenotypic data was obtained from (Yanarella et al., 2023a) and is available from: https://datacommons.cyverse.org/browse/iplant/home/shared/commons_repo/curated/Carolyn_Lawrence_Dill_Maize_WiDiv_Summer_2021_Dataset_June_2023. Genotypic data was obtained from (Mural et al., 2022b,a) and can be accessed from: https://figshare.com/articles/dataset/Maize_WiDiv_SAM_1051Genotype_vcf_gz_genotype_file/19175888/1. Gene Ontology data was obtained from (Wimalanathan and Lawrence-Dill, 2017) and is available from https://datacommons.cyverse.org/browse/iplant/home/shared/commons_repo/curated/Carolyn_Lawrence-Dill_maize-GAMER_maize.B73_RefGen_v4_Zm00001d.2_Oct_2017.r1.

## 7 ACKNOWLEDGMENTS

We acknowledge the High-Performance Computing (HPC) facility at Iowa State University for assisting this research through computing resources and technical support. We appreciate discussions with Toni Kazic about the preliminary conceptualizations for this research project and making audio data available to assist with experimental design. We appreciate discussions with Qi Li regarding the binning methods and Kris De Brabanter for discussions about the statistics underlying GWAS. We thank Jianming Yu for helpful discussions about genomic datasets and GWAS results. We thank Brian Dilkes, Rajdeep Khangura, and Amanpreet Kaur for providing genotypic data, guidance, and examples of data processing techniques for association studies. We thank Tyler Foster and Yu-Ru Chen for enlightening discussions about GWAS methods and tools. We thank reviewers of the submitted manuscript for helpful guidance to improve the paper.

## 10 AUTHOR CONTRIBUTIONS

CFY and CJLD conceived the idea for this project. CFY performed the analyses and drafted the initial version of the manuscript. CFY and LF evaluated the GO term analysis. All authors have read, edited, and approved the manuscript.

## REFERENCES

Abadi, M., Barham, P., Chen, J., Chen, Z., Davis, A., Dean, J., Devin, M., Ghemawat, S., Irving, G., Isard, M., et al. (2016). Tensorflow: A system for large-scale machine learning. In 12th USENIX Symposium on Operating Systems Design and Implementation (OSDI ‘16), pages 265–283.

Andorf, C. M., Cannon, E. K., Portwood, J. L., Gardiner, J. M., Harper, L. C., Schaeffer, M. L., Braun, B. L., Campbell, D. A., Vinnakota, A. G., Sribalusu, V. V., et al. (2016). Maizegdb update: new tools, data and interface for the maize model organism database. Nucleic Acids Research, 44(D1):D1195–D1201.

Austin, D. F. and Lee, M. (1996). Genetic resolution and verification of quantitative trait loci for flowering and plant height with recombinant inbred lines of maize. Genome, 39(5):957–968.

Azodi, C. B., Pardo, J., VanBuren, R., de los Campos, G.x, and Shiu, S.-H. (2020). Transcriptome-based prediction of complex traits in maize. The Plant Cell, 32(1):139–151.

Bai, W., Zhang, H., Zhang, Z., Teng, F., Wang, L., Tao, Y., and Zheng, Y. (2009). The evidence for non-additive effect as the main genetic component of plant height and ear height in maize using introgression line populations. Plant Breeding.

Bates, D., Mächler, M., Bolker, B., and Walker, S. (2015). Fitting linear mixed-effects models using lme4. Journal of Statistical Software, 67(1):1–48.

Blakeslee, J. J., Peer, W. A., and Murphy, A. S. (2005). Auxin transport. Current Opinion in Plant Biology, 8(5):494–500.

Bradbury, P. J., Zhang, Z., Kroon, D. E., Casstevens, T. M., Ramdoss, Y., and Buckler, E. S. (2007). TASSEL: software for association mapping of complex traits in diverse samples. Bioinformatics, 23(19):2633–2635.

Braun, I. R. and Lawrence-Dill, C. J. (2020). Automated methods enable direct computation on phenotypic descriptions for novel candidate gene prediction. Frontiers in Plant Science, 10.

Braun, I. R., Yanarella, C. F., and Lawrence-Dill, C. J. (2020). Computing on phenotypic descriptions for candidate gene discovery and crop improvement. Plant Phenomics, 2020.

Braun, I. R., Yanarella, C. F., Rajeswari, J. P. D., Bassham, D. C., and Lawrence-Dill, C. J. (2021). The Case for Retaining Natural Language Descriptions of Phenotypes in Plant Databases and a Web Application as Proof of Concept. bioRxiv.

Brooks, L., Strable, J., Zhang, X., Ohtsu, K., Zhou, R., Sarkar, A., Hargreaves, S., Elshire, R. J., Eudy, D., Pawlowska, T., Ware, D., Janick-Buckner, D., Buckner, B., Timmermans, M. C. P., Schnable, P. S., Nettleton, D., and Scanlon, M. J. (2009). Microdissection of shoot meristem functional domains. PLoS Genetics, 5(5):e1000476.

Carlson, M. (2023). GO.db: A set of annotation maps describing the entire Gene Ontology. R package version 3.17.0.

Danecek, P., Auton, A., Abecasis, G., Albers, C. A., Banks, E., DePristo, M. A., Handsaker, R. E., Lunter, G., Marth, G. T., Sherry, S. T., McVean, G., and and, R. D. (2011). The variant call format and VCFtools. Bioinformatics, 27(15):2156–2158.

Fattel, L., Psaroudakis, D., Yanarella, C. F., Chiteri, K. O., Dostalik, H. A., Joshi, P., Starr, D. C., Vu, H., Wimalanathan, K., and Lawrence-Dill, C. J. (2022). Standardized genome-wide function prediction enables comparative functional genomics: a new application area for gene ontologies in plants. GigaScience, 11.

Fornalé, S., Rencoret, J., Garcia-Calvo, L., Capellades, M., Encina, A., Santiago, R., Rigau, J., Gutiérrez, A., Del Río, J.-C., and Caparros-Ruiz, D. (2015). Cell wall modifications triggered by the down-regulation of coumarate 3-hydroxylase-1 in maize. Plant Science, 236:272–282.

Fu, J., Zhang, D.-F., Liu, Y.-H., Ying, S., Shi, Y.-S., Song, Y.-C., Li, Y., and Wang, T.-Y. (2012). Isolation and characterization of maize pmp3 genes involved in salt stress tolerance. PloS one, 7(2):e31101.

Gallavotti, A. (2013). The role of auxin in shaping shoot architecture. Journal of Experimental Botany, 64(9):2593–2608.

Galli, M., Liu, Q., Moss, B. L., Malcomber, S., Li, W., Gaines, C., Federici, S., Roshkovan, J., Meeley, R., Nemhauser, J. L., et al. (2015). Auxin signaling modules regulate maize inflorescence architecture. Proceedings of the National Academy of Sciences, 112(43):13372–13377.

Geisler, M. and Murphy, A. S. (2005). The ABC of auxin transport: The role of p-glycoproteins in plant development. FEBS Letters, 580(4):1094–1102.

German, C. A., Sinsheimer, J. S., Klimentidis, Y. C., Zhou, H., and Zhou, J. J. (2019). Ordered multinomial regression for genetic association analysis of ordinal phenotypes at biobank scale. Genetic Epidemiology, 44(3):248–260.

Goode, K. and Rey, K. (2022). ggResidpanel: Panels and Interactive Versions of Diagnostic Plots using ‘ggplot2’. R package version 0.3.0.

Hamazaki, K. and Iwata, H. (2020). RAINBOW: Haplotype-based genome-wide association study using a novel SNP-set method. PLOS Computational Biology, 16(2):e1007663.

Hansey, C. N., Johnson, J. M., Sekhon, R. S., Kaeppler, S. M., and de Leon, N. (2011). Genetic diversity of a maize association population with restricted phenology. Crop Science, 51(2):704–715.

Hartwig, T., Chuck, G. S., Fujioka, S., Klempien, A., Weizbauer, R., Potluri, D. P. V., Choe, S., Johal, G. S., and Schulz, B. (2011). Brassinosteroid control of sex determination in maize. Proceedings of the National Academy of Sciences, 108(49):19814–19819.

Hirsch, C. N., Foerster, J. M., Johnson, J. M., Sekhon, R. S., Muttoni, G., Vaillancourt, B., Peñagaricano, F., Lindquist, E., Pedraza, M. A., Barry, K., de Leon, N., Kaeppler, S. M., and Buell, C. R. (2014). Insights into the maize pan-genome and pan-transcriptome . The Plant Cell, 26(1):121–135.

Honnibal, M. and Montani, I. (2023). spaCy v3.5.1 spancat for multi-class labeling, fixes for textcat+transformers and more. To appear.

Jansson, S. (1994). The light-harvesting chlorophyll ab-binding proteins. Biochimica et Biophysica Acta (BBA) - Bioenergetics, 1184(1):1–19.

Kat IP Pty Ltd (2008). WordHippo.

Kazic, T. (2020). Chloe: Flexible, efficient data provenance and management. bioRxiv.

Khanna, R., Shen, Y., Toledo-Ortiz, G., Kikis, E. A., Johannesson, H., Hwang, Y.-S., and Quail, P. H. (2006). Functional profiling reveals that only a small number of phytochrome-regulated early-response genes in arabidopsis are necessary for optimal deetiolation. The Plant Cell, 18(9):2157–2171.

Khavkin, E. and Coe, E. H. (1997). Mapped genomic locations for developmental functions and qtls reflect concerted groups in maize (zea mays l.). Theoretical and Applied Genetics, 95:343–352.

Koroleva, A., Kamath, S., and Paroubek, P. (2019). Measuring semantic similarity of clinical trial outcomes using deep pre-trained language representations. Journal of Biomedical Informatics, 100:100058.

Lawit, S. J., Wych, H. M., Xu, D., Kundu, S., and Tomes, D. T. (2010). Maize DELLA proteins dwarf plant8 and dwarf plant9 as modulators of plant development. Plant and Cell Physiology, 51(11):1854–1868.

Lee, J., Yoon, W., Kim, S., Kim, D., Kim, S., So, C. H., and Kang, J. (2019). BioBERT: a pre-trained biomedical language representation model for biomedical text mining. Bioinformatics, 36(4):1234–1240.

Lenth, R. V. (2023). emmeans: Estimated Marginal Means, aka Least-Squares Means. R package version 1.8.7.

Li, H., Wang, L., Liu, M., Dong, Z., Li, Q., Fei, S., Xiang, H., Liu, B., and Jin, W. (2020). Maize plant architecture is regulated by the ethylene biosynthetic gene ZmACS7. Plant Physiology, 183(3):1184–1199.

Lipka, A. E., Tian, F., Wang, Q., Peiffer, J., Li, M., Bradbury, P. J., Gore, M. A., Buckler, E. S., and Zhang, Z. (2012). GAPIT: genome association and prediction integrated tool. Bioinformatics, 28(18):2397–2399.

Liu, X., Huang, M., Fan, B., Buckler, E. S., and Zhang, Z. (2016). Iterative usage of fixed and random effect models for powerful and efficient genome-wide association studies. PLOS Genetics, 12(2):e1005767.

Mazaheri, M., Heckwolf, M., Vaillancourt, B., Gage, J. L., Burdo, B., Heckwolf, S., Barry, K., Lipzen, A., Ribeiro, C. B., Kono, T. J. Y., Kaeppler, H. F., Spalding, E. P., Hirsch, C. N., Buell, C. R., de Leon, N., and Kaeppler, S. M. (2019). Genome-wide association analysis of stalk biomass and anatomical traits in maize. BMC Plant Biology, 19(1).

Mensio, M. (2023). Martinomensio/spacy-universal-sentence-encoder: Google use (universal sentence encoder) for spacy.

Merriam-Webster (2023). Merriam-Webster Online Thesaurus.

Multani, D. S., Briggs, S. P., Chamberlin, M. A., Blakeslee, J. J., Murphy, A. S., and Johal, G. S. (2003). Loss of an MDR transporter in compact stalks of maize br2 and sorghum dw3 mutants. Science, 302(5642):81–84.

Mungall, C. J., Gkoutos, G. V., Smith, C. L., Haendel, M. A., Lewis, S. E., and Ashburner, M. (2010). Integrating phenotype ontologies across multiple species. Genome Biology, 11(1):R2.

Mural, R., Sun, G., Grzybowski, M., Tross, M. C., Jin, H., Smith, C., Newton, L., Thompson, A. M., Sigmon, B., and Schnable, J. C. (2022a). Maize WiDiv SAM 1051Genotype.vcf.gz genotype file.

Mural, R. V., Sun, G., Grzybowski, M., Tross, M. C., Jin, H., Smith, C., Newton, L., Andorf, C. M., Woodhouse, M. R., Thompson, A. M., Sigmon, B., and Schnable, J. C. (2022b). Association mapping across a multitude of traits collected in diverse environments in maize. GigaScience, 11.

Oellrich, A., Walls, R. L., Cannon, E. K., Cannon, S. B., Cooper, L., Gardiner, J., Gkoutos, G. V., Harper, L., He, M., Hoehndorf, R., Jaiswal, P., Kalberer, S. R., Lloyd, J. P., Meinke, D., Menda, N., Moore, L., Nelson, R. T., Pujar, A., Lawrence, C. J., and Huala, E. (2015). An ontology approach to comparative phenomics in plants. Plant Methods, 11(1).

Peiffer, J. A., Romay, M. C., Gore, M. A., Flint-Garcia, S. A., Zhang, Z., Millard, M. J., Gardner, C. A. C., McMullen, M. D., Holland, J. B., Bradbury, P. J., and Buckler, E. S. (2014). The genetic architecture of maize height. Genetics, 196(4):1337–1356.

R Core Team (2023). R: A Language and Environment for Statistical Computing. R Foundation for Statistical Computing, Vienna, Austria.

Řehůřek, R. and Sojka, P. (2010). Software Framework for Topic Modelling with Large Corpora. In Proceedings of the LREC 2010 Workshop on New Challenges for NLP Frameworks, pages 45–50, Valletta, Malta. ELRA.

Reiser, L., Subramaniam, S., Zhang, P., and Berardini, T. (2022). Using the arabidopsis information resource (tair) to find information about arabidopsis genes. Current protocols, 2(10):e574.

Salvi, S., Corneti, S., Bellotti, M., Carraro, N., Sanguineti, M. C., Castelletti, S., and Tuberosa, R. (2011). Genetic dissection of maize phenology using an intraspecific introgression library. BMC Plant Biology, 11(1):4.

Sarić, R., Nguyen, V. D., Burge, T., Berkowitz, O., Trtílek, M., Whelan, J., Lewsey, M. G., and Čustović, E. (2022). Applications of hyperspectral imaging in plant phenotyping. Trends in Plant Science, 27(3):301–315.

Stein, L. D. (2013). Using GBrowse 2.0 to visualize and share next-generation sequence data. Briefings in Bioinformatics, 14(2):162–171.

Sterck, L. (2021). Calculate and draw custom Venn diagrams.

Tang, Y., Liu, X., Wang, J., Li, M., Wang, Q., Tian, F., Su, Z., Pan, Y., Liu, D., Lipka, A. E., Buckler, E. S., and Zhang, Z. (2016). GAPIT version 2: An enhanced integrated tool for genomic association and prediction. The Plant Genome, 9(2).

Teng, F., Zhai, L., Liu, R., Bai, W., Wang, L., Huo, D., Tao, Y., Zheng, Y., and Zhang, Z. (2012). ZmGA3ox2, a candidate gene for a major QTL, qPH3.1, for plant height in maize. The Plant Journal, 73(3):405–416.

Van Rossum, G. and Drake, F. L. (2009). Python 3 Reference Manual. CreateSpace, Scotts Valley, CA.

Venables, W. N. and Ripley, B. D. (2002). Modern Applied Statistics with S. Springer, New York, fourth edition. ISBN 0-387-95457-0.

Vendramin, S., Huang, J., Crisp, P. A., Madzima, T. F., and McGinnis, K. M. (2020). Epigenetic regulation of aba-induced transcriptional responses in maize. G3: Genes, Genomes, Genetics, 10(5):1727–1743.

Wallace, J. G., Zhang, X., Beyene, Y., Semagn, K., Olsen, M., Prasanna, B. M., and Buckler, E. S. (2016). Genome-wide association for plant height and flowering time across 15 tropical maize populations under managed drought stress and well-watered conditions in sub-saharan africa. Crop Science, 56(5):2365–2378.

Wang, J. and Zhang, Z. (2021). GAPIT version 3: Boosting power and accuracy for genomic association and prediction. Genomics, Proteomics & Bioinformatics, 19(4):629–640.

Wang, Q., Song, S., Lu, X., Wang, Y., Chen, Y., Wu, X., Tan, L., and Chai, G. (2022). Hormone regulation of ccch zinc finger proteins in plants. International Journal of Molecular Sciences, 23(22):14288.

Weng, J., Xie, C., Hao, Z., Wang, J., Liu, C., Li, M., Zhang, D., Bai, L., Zhang, S., and Li, X. (2011). Genome-wide association study identifies candidate genes that affect plant height in chinese elite maize (Zea mays L.) inbred lines. PLoS ONE, 6(12):e29229.

Wickham, H. (2016). ggplot2: Elegant Graphics for Data Analysis. Springer-Verlag New York.

Wimalanathan, K., Friedberg, I., Andorf, C. M., and Lawrence-Dill, C. J. (2018). Maize G. annotation—methods, evaluation, and review (maize-GAMER). Plant Direct, 2(4).

Wimalanathan, K. and Lawrence-Dill, C. (2017). maize-GAMER Annotations for maize B73 RefGen V4 Zm00001d.2.

Winkler, R. G. and Helentjaris, T. (1995). The maize dwarf3 gene encodes a cytochrome p450-mediated early step in gibberellin biosynthesis. The Plant Cell, 7(8):1307–1317.

Woodhouse, M. R., Cannon, E. K., Portwood, J. L., Harper, L. C., Gardiner, J. M., Schaeffer, M. L., and Andorf, C. M. (2021). A pan-genomic approach to genome databases using maize as a model system. BMC Plant Biology, 21(1).

Wu, A.-M., Rihouey, C., Seveno, M., Hörnblad, E., Singh, S. K., Matsunaga, T., Ishii, T., Lerouge, P., and Marchant, A. (2009). The arabidopsis IRX10 and IRX10-LIKE glycosyltransferases are critical for glucuronoxylan biosynthesis during secondary cell wall formation. The Plant Journal, 57(4):718–731.

Xiao, Q., Bai, X., Zhang, C., and He, Y. (2022). Advanced high-throughput plant phenotyping techniques for genome-wide association studies: A review. Journal of Advanced Research, 35:215–230.

Xu, C., Sato, Y., Yamazaki, M., Brasser, M., Barbour, M. A., Bascompte, J., and Shimizu, K. K. (2023). Genome-wide association study of aphid abundance highlights a locus affecting plant growth and flowering in arabidopsis thaliana. Royal Society Open Science, 10(8):230399.

Yanarella, C. F., Fattel, L., Kristmundsdóttir Á Ý, Lopez, M. D., Edwards, J. W., Campbell, D. A., Abel, C. A., and Lawrence-Dill, C. J. (2023a). Carolyn Lawrence Dill Maize WiDiv Summer 2021 Dataset June 2023.

Yanarella, C. F., Fattel, L., Kristmundsdóttir Á Ý, Lopez, M. D., Edwards, J. W., Campbell, D. A., Abel, C. A., and Lawrence-Dill, C. J. (2024). Wisconsin diversity panel phenotypes: spoken descriptions of plants and supporting data. BMC Research Notes, 17(1).

Yanarella, C. F., Fattel, L., and Lawrence-Dill, C. J. (2023b). Carolyn Lawrence Dill Maize WiDiv Association Studies Dataset September 2023.

Yang, W., Feng, H., Zhang, X., Zhang, J., Doonan, J. H., Batchelor, W. D., Xiong, L., and Yan, J. (2020). Crop phenomics and high-throughput phenotyping: Past decades, current challenges, and future perspectives. Molecular Plant, 13(2):187–214.

Yao, L., van de Zedde, R., and Kowalchuk, G. (2021). Recent developments and potential of robotics in plant eco-phenotyping. Emerging Topics in Life Sciences, 5(2):289–300.

Yu, J., Pressoir, G., Briggs, W. H., Bi, I. V., Yamasaki, M., Doebley, J. F., McMullen, M. D., Gaut, B. S., Nielsen, D. M., Holland, J. B., Kresovich, S., and Buckler, E. S. (2005). A unified mixed-model method for association mapping that accounts for multiple levels of relatedness. Nature Genetics, 38(2):203–208.

Zhang, C., Dong, S.-S., Xu, J.-Y., He, W.-M., and Yang, T.-L. (2018). PopLDdecay: a fast and effective tool for linkage disequilibrium decay analysis based on variant call format files. Bioinformatics, 35(10):1786–1788.

