## Supplementary Figure 1 for "GWAS from Spoken Phenotypic Descriptions: A Proof of Concept from Maize Field Studies"

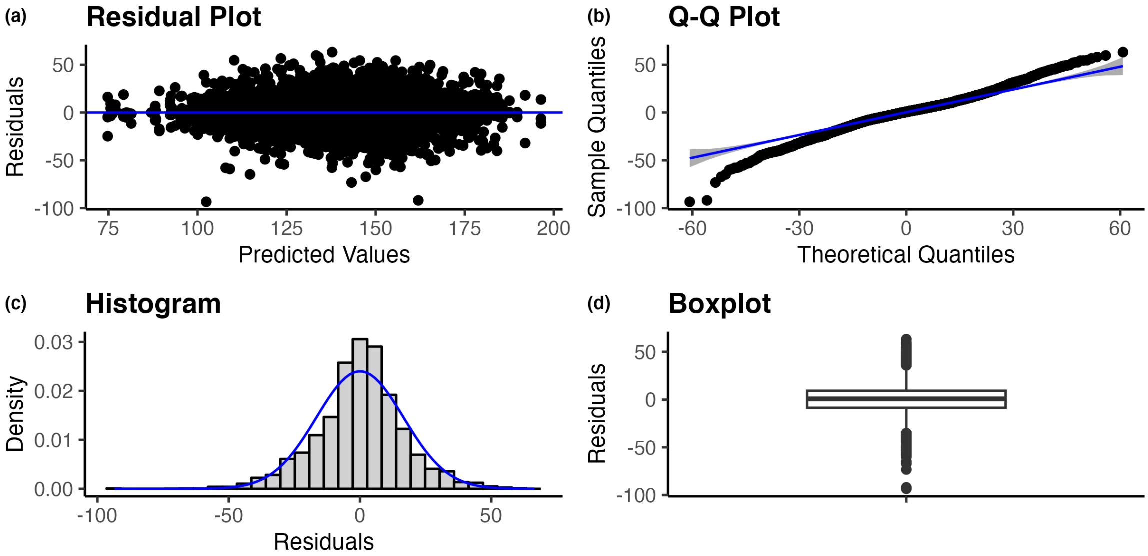


Supplementary Figure 1 Diagnostic plots for the linear regression model used for BLUEs estimation for measured plant height data. (a) Residual plot of the residuals versus the predicted values of the model, (b) Q−Q Plot of sample vs theoretical quantiles, (c) Histogram of residuals with normal density curve overlayed, (d) Boxplot of residuals.
