## Supplementary Figure 3 for "GWAS from Spoken Phenotypic Descriptions: A Proof of Concept from Maize Field Studies"

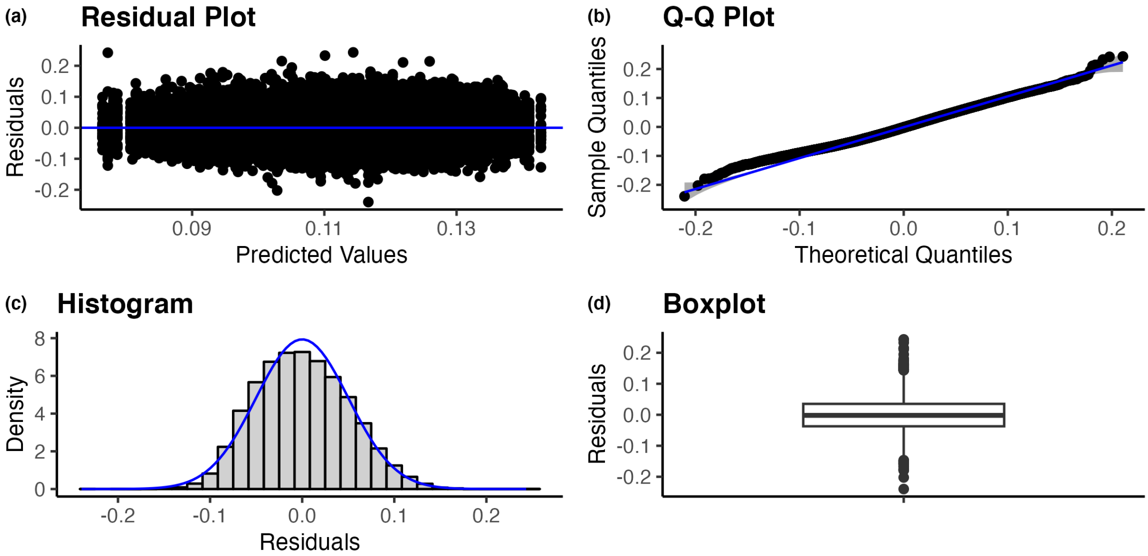


Supplementary Figure 3 Diagnostic plots for the linear regression model used for BLUEs estimation for tall query semantic similarity data. (a) Residual plot of the residuals versus the predicted values of the model, (b) Q−Q Plot of sample vs theoretical quantiles, (c) Histogram of residuals with normal density curve overlayed, (d) Boxplot of residuals.
