## Supplementary Figure 7 for "GWAS from Spoken Phenotypic Descriptions: A Proof of Concept from Maize Field Studies"

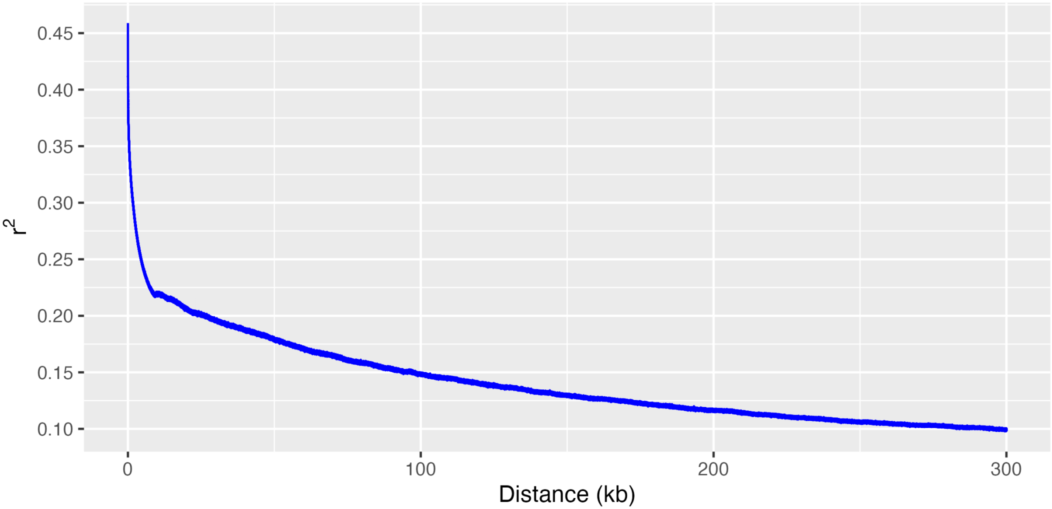


Supplementary Figure 7 LD decay curve using Mural et al. data taxa intersecting with Yanarella et al. taxa. Blue line represents the plot of r^2 vs distance in kb.
