## Supplementary Figure 8 for "GWAS from Spoken Phenotypic Descriptions: A Proof of Concept from Maize Field Studies"

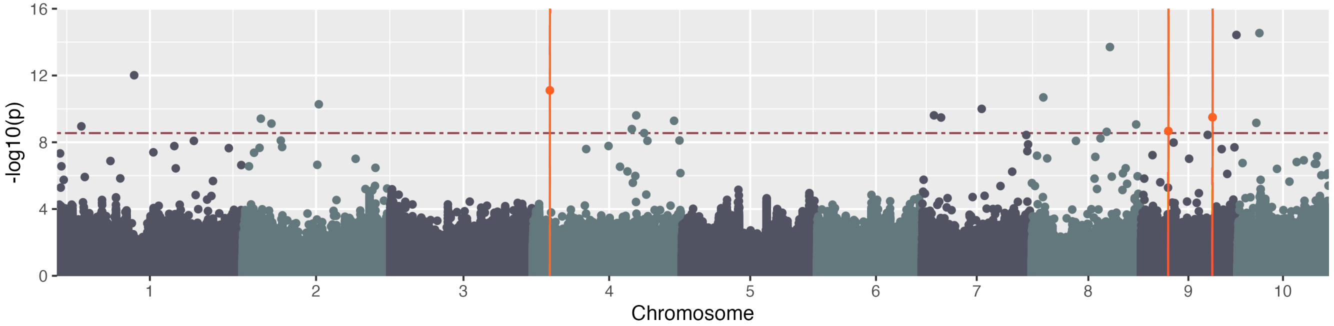


Supplementary Figure 8 Manually binned spoken height - phenotypic data. Manhattan plot generated using GAPIT and FarmCPU using manually binned phrases and the multinomial regression using Mural et al. genotypic data. The red dashed line indicates the Bonferroni threshold ($-\log_{10}(p) = 8.55$), orange points indicate identified SNPs with known plant height genes within +/− 300 kb, and orange vertical lines indicate positions +/− 300 kb identified SNPs with known plant height genes.
